# Profiling of RNA 8-oxoG marks in *Escherichia coli* identifies critical intrinsic characteristics that contribute to 8-oxoG accumulation in bacteria

**DOI:** 10.1101/2025.10.14.682464

**Authors:** Matthew R. Burroughs, Blanca I. Quiñones-Diaz, Lydia M. Contreras

**Author notes:** Address correspondence to Dr. Lydia Contreras. Email Addresses: Matthew R. Burroughs, Blanca I. Quiñones-Diaz, Lydia M. Contreras.

## Abstract

Reactive oxygen species (ROS) are environmentally ubiquitous and known to have pervasive impacts on cellular homeostasis. RNA is vulnerable to oxidative chemical alterations from a variety of endogenous and exogenous sources. The most common chemical modification resulting from ROS exposure to RNA is 8-oxo-7,8-dihydroguanine (8-oxoG)—an oxidized form of the canonical guanine (G) nucleobase. While 8-oxoG modifications are known to impact mRNA processing, understanding the broader biological impact of 8-oxoG requires knowledge of how these modifications accumulate. In this work, we assessed the disparate enrichment of 8-oxoG modifications within RNAs in the *E. coli* transcriptome using an RNA Immunoprecipitation Sequencing technique with a high-affinity 8-oxoG antibody (8-oxoG-RIP-Seq). Our investigation of the RNA 8-oxoG enrichment landscape uncovered several intrinsic RNA characteristics that correlate with 8-oxoG enrichment. These findings suggest intrinsic characteristics of RNA, most notably relative abundance, CDS length, and G nucleotide composition, significantly influence RNA 8-oxoG accumulation. We harnessed these intrinsic characteristics to construct a simple multiple linear regression model that predicts RNA 8-oxoG accumulation, which we validated in *E. coli*. This model was subsequently applied to predict 8-oxoG enriched RNA species in four other bacterial species spanning a wide range of oxidative stress tolerances; these predictions suggest that 8-oxoG accumulation is largely species dependent, with limited overlap in RNAs and functional pathways that are more susceptible to elevated levels of 8-oxoG accumulation. Overall, these findings better inform understanding of RNA 8-oxoG patterns in bacteria and have broader impacts towards advancing knowledge of the connection between RNA oxidation and cellular homeostasis.

## Introduction

Bacteria frequently encounter reactive oxygen species (ROS) in the environment from a variety of sources (1) including UV-mediated photochemistry in marine and freshwater surfaces (2,3), oxic-anoxic interfaces present in the soil microbiome (4), and phagocytosis processes within eukaryotic host organisms (5). Concurrently, endogenous production of ROS is triggered through fundamental cellular redox processes (6). Oxidative stress manifests from the overabundance of ROS in the cell, overwhelming host defense systems. Bacteria have evolved a suite of mechanisms to mitigate the proliferation of ROS such as redox-sensitive transcription factor activation/repression cascades (7,8), DNA damage response (DDR) pathways (9), ROS scavenging enzymes (10), and other enzymes capable of degrading oxidized nucleotides within the free nucleotide pool (11). As an illustrative example, in many bacteria, the well-characterized OxyR transcription factor is sensitive to H_2_O_2_-mediated oxidation of cysteine residue 199 (7). This oxidation event triggers a conformational shift in the protein and biases its DNA interactome, ultimately leading to the transcriptional activation of a defined suite of antioxidant genes (8). Tolerance to oxidative stress varies widely across bacteria, with some species such as *D. radiodurans* capable of withstanding immense oxidative stress insult (12) while others including *S. oneidensis* are far more sensitive to ROS exposure (13). This observed discrepancy, e.g. in the case of radiation tolerance, has been largely attributed to the relative ratios of intracellular manganese to iron (14,15), with these elements acting as repressors or drivers of ROS potency, respectively. However, additional factors (e.g. antioxidant protein activities, genomic compositions (16)) also likely contribute to the observed range of oxidative stress tolerance across bacterial species.

Oxidative stress is known to chemically alter biomolecules in the cell including proteins, lipids, and nucleic acids (17,18). RNA is uniquely sensitive to oxidative damage (19–21), due in part to its single-stranded primary structure, high intracellular abundance, lack of nucleobase repair machinery, and lower average protein occupancy compared to DNA (22). Within RNA, the canonical guanine (G) nucleobase is the most frequent target of oxidative damage—a result of its low redox potential relative to the other canonical nucleobases (23). Oxidative damage to G can propagate a plethora of chemically modified derivatives; however, 8-oxo-7,8-dihydroguanine (8-oxoG), a two electron oxidation product, is the most prevalent RNA modification to arise from oxidation of this nucleobase (24). Other RNA modifications are believed to play an indirect role in response to oxidative stress induced damage, namely through cursory involvement in the DNA damage response (25).

RNA oxidation accumulates disparately across the transcriptome, with certain RNAs being naturally enriched or depleted in 8-oxoG modifications (26–31). Studies which have investigated 8-oxoG modifications in DNA and RNA identified certain sequence features that appear to partially dictate the uneven distribution of these modifications (26,32–36). Prior work in DNA has shown that 8-oxoG modifications are more common within GC-rich promoter regions upstream of gene coding sequences (32,33). Additional DNA studies have found enrichment of 8-oxoG lesions at genomic G-quadruplex (G4) formation sites within gene promoter sequences, wherein subsequent recruitment of DNA base excision repair (BER) machinery ultimately drives elevated transcription of downstream genes (34,35,37). More recently, a transcriptome-wide analysis of 8-oxoG modifications in human lung BEAS-2B cells identified that 8-oxoG modifications are largely enriched within RNAs that are elevated in length and G nucleotide composition (26). Similarly, a study in HEK293T cells revealed elevated G oxidation events in the GC-rich tentacles of 28S ribosomal RNA sequences (36). In total, these prior investigations strongly suggest that many RNA oxidation events are a function of defined intrinsic RNA variables.

The 8-oxoG modification has garnered increased interest as a human health biomarker, since oxidative stress is a widespread hallmark of various disease pathologies and aging (reviewed in (38)). Initial mechanistic studies have indicated that 8-oxoG impacts RNA fate predominately through ribosomal stalling and subsequent triggering of no-go decay (NGD) surveillance pathways (39–41). Additionally, the mutagenic base-pairing potential of this modification in the *syn* conformation to canonical adenine (A) residues has been shown to impact tRNA anticodon alignment during translation (42). Unlike other well-studied RNA modifications (e.g. m^6^A, m^5^C, Ψ), 8-oxoG modifications are not known to be catalytically installed by specific writer enzymes; instead, these modifications are believed to arise solely from direct oxidation of RNA by neighboring ROS and are thus classified as a non-enzymatic modification (NECM) (43). Reader proteins which selectively recognize and bind to 8-oxoG modified RNAs over unmodified RNAs have been identified (44), although understanding the broader implications of these selective binding interactions continues to be an area of active research. For example, *E. coli* polynucleotide phosphorylase (PNPase), an exoribonuclease, has been shown to selectively bind to oxidized RNAs over non-oxidized RNAs (45,46) and, interestingly, to stall at 8-oxoG sites in RNA in vitro (47). While the preference of this exoribonuclease for oxidized RNA templates suggests an evolved mechanism for targeted, expedited RNA clearance, the larger systems-level consequence of this observed stalling phenomenon remains unknown. Overall, multifaceted downstream implications of this unique RNA modification warrant additional transcriptome-wide studies.

Largely owing to the intimate link between RNA oxidation and human disease pathologies, an overwhelming majority of the studies performed to date which have investigated 8-oxoG RNA modification landscapes have been performed in eukaryotic cell models which span mRNA (26–29,31), rRNA (36), miRNA (48–50), lncRNA (30), and circRNA (51,52) subclasses. Despite increasing work investigating RNA oxidation in eukaryotic organisms, a comparatively small number of studies have investigated the landscape of RNA 8-oxoG modifications in bacteria. To date, the work investigating RNA 8-oxoG landscapes in bacteria has been limited to ribosomal RNAs (53). Given the importance of understanding basic principles of RNA biology, particularly as it relates to oxidative stress responses in bacteria, we herein investigate the disparate accumulation of 8-oxoG modifications in the model bacterium *E. coli*. We first performed a transcriptome-wide investigation of the landscape of 8-oxoG modifications in ribosomally-depleted RNA under basal and oxidatively stressed growth conditions through 8-oxoG RNA Immunoprecipitation Sequencing (8-oxoG-RIP-Seq). After identifying RNAs that are relatively enriched or depleted in 8-oxoG modifications, we confirmed several of the 8-oxoG enriched RNAs by performing the analogous Chemical Labeling of RNA targeting 8-oxoG modifications and Sequencing (ChLoRox-Seq (26)) under the same experimental conditions. Upon analysis of specific biological pathways that are enriched in RNAs with elevated 8-oxoG accumulation, we observed pathways related to various transport processes. Our closer investigation of 8-oxoG enriched and depleted RNAs uncovered that several intrinsic RNA characteristics are correlated with RNA 8-oxoG enrichment; these properties notably include relative cellular abundance, CDS length, and G nucleotide composition. Lastly, we harnessed observed patterns in intrinsic RNA characteristics to construct a multiple linear regression machine learning model that predicts mRNA 8-oxoG accumulation. We subsequently applied this model to predict 8-oxoG enriched RNA species in four other bacterial species spanning a wide range of oxidative stress tolerances; these predictions suggest that 8-oxoG accumulation is largely species dependent. In this way, we observe that RNA 8-oxoG modifications accumulate within different genes and biological pathways in a manner that is unique to each bacterial species, and that these differences might partially explain the range of oxidative stress tolerance observed across bacteria.

## Results

### 8-oxoG-RIP-Seq captures RNA transcripts enriched and depleted in 8-oxoG modifications in E. coli across oxidatively stressed and unstressed cell states

We began by profiling the RNA 8-oxoG modification landscape in the model organism *E. coli* using an adapted 8-oxoG-RIP-Seq method (Fig. 1A) (27,28). This method utilizes specific binding of a commercially available antibody to the 8-oxoG modification, enabling subsequent protein G-based affinity enrichment and next generation sequencing (NGS) to identify RNAs with elevated 8-oxoG accumulation. We assayed cells grown to exponential phase (OD ∼ 0.5-0.7) that were subsequently subjected to either 1 mM H_2_O_2_ (H_2_O_2_ Treated) or PBS (Untreated) treatment for 20 minutes at 37°C. These sublethal H_2_O_2_ exposure conditions were sufficient to induce elevated expression of known oxidative stress response genes (e.g. *oxyS*, *recA*, *sodA*, etc.) (Supplemental Fig. S1A, Supplemental Table S1) without leading to significant increases in bulk 8-oxoG levels (Supplemental Fig. S1B). Through 8-oxoG-RIP-Seq analysis, we identified subpopulations of RNAs with elevated levels of 8-oxoG accumulation (8-oxoG enriched, log2FoldChange(IP/Input) > 1, p_adj_ < 0.05) and reduced levels of 8-oxoG accumulation (8-oxoG depleted, log2FoldChange(IP/Input) < -1, p_adj_ < 0.05) in both Untreated (Fig. 1B) and H_2_O_2_ Treated (Fig. 1C) experimental conditions. In total, we found 459 RNAs enriched in 8-oxoG under Untreated conditions and 383 such RNAs under H_2_O_2_ Treated conditions (Fig. 1D). Conversely, we identified 321 RNAs depleted in 8-oxoG under Untreated conditions and 186 such RNAs under H_2_O_2_ Treated conditions (Fig. 1E). Many of the RNA transcripts were similarly enriched in 8-oxoG across both conditions (Fig. 1F), suggesting a degree of stress-independence for RNAs that are enriched in 8-oxoG under basal conditions. Approximately 50% of 8-oxoG enriched RNAs (283) were shared across both tested conditions (Fig. 1D). Similarly, roughly 40% of 8-oxoG depleted RNAs (149) were conserved across conditions (Fig. 1E). We further evaluated RNAs which appear to undergo differential 8-oxoG enrichment under H_2_O_2_ exposure conditions (|log2FoldChange(IP/Input)_H2O2_ – log2FoldChange(IP/Input)_Untreated_| > 1.5). Interestingly, we saw the emergence of several regulatory small RNAs (sRNAs) (e.g. *fecD*, *fnrS*, *dsrA*) within this differentially oxidized population (Supplemental Fig. S2).

**Figure 1.**
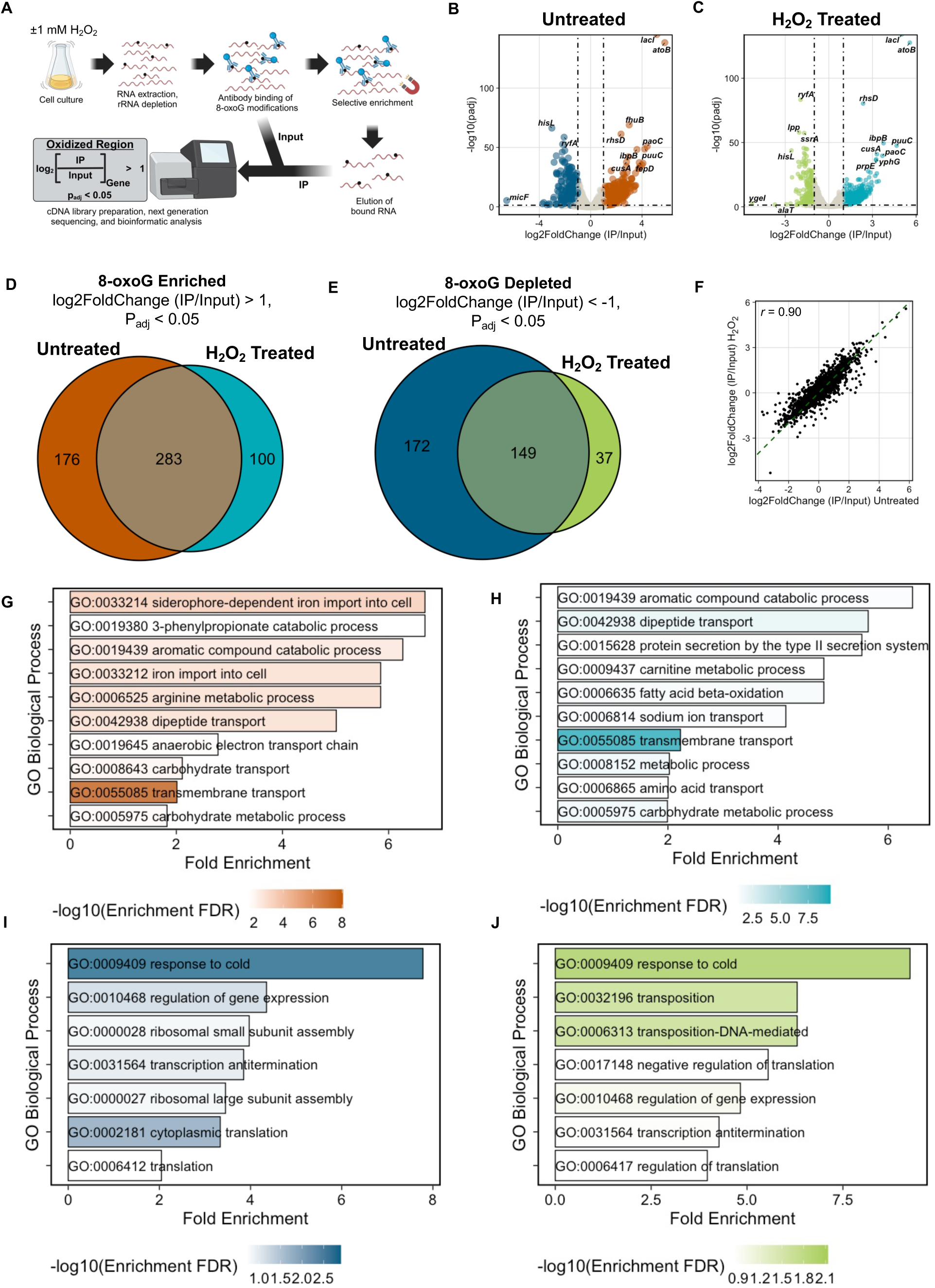
8-oxoG-RIP-Seq captures transcripts enriched in 8-oxoG modifications in *E. coli* across Untreated and H_2_O_2_ Treated growth conditions. (A) Diagram of 8-oxoG-RIP-Seq pipeline in *E. coli*. (B) Volcano plot of 8-oxoG enriched (orange) and 8-oxoG depleted (blue) RNAs in Untreated growth condition. (C) Volcano plot of 8-oxoG enriched (cyan) and 8-oxoG depleted (green) RNAs in H_2_O_2_ Treated growth condition. (D-E) Euler plots showing overlapping 8-oxoG enriched (E) and 8-oxoG depleted (F) genes across growth conditions. (F) Correlation plot comparing log2FoldChange(IP/Input) 8-oxoG enrichment values across growth conditions. *r* = Pearson’s correlation coefficient. Dashed line represents y = x. (G-J) GO Biological Process gene overrepresentation analysis plots of pathways elevated in (G) 8-oxoG enriched RNAs under Untreated growth condition, (H) 8-oxoG enriched RNAs under H_2_O_2_ Treated growth condition, (I) 8-oxoG depleted RNAs under Untreated growth condition, and (J) 8-oxoG depleted RNAs under H_2_O_2_ Treated growth condition.

We next performed GO pathway over representation analysis to determine whether RNAs within specific biological process pathways were elevated in 8-oxoG enriched or depleted RNA species under each experimental condition (Fig. 1G-J). The transmembrane transport pathway (GO:0055085) was identified to be elevated in 8-oxoG enriched RNAs relative to background genes for both experimental conditions (Fig. 1G-H). Conversely, response to cold (GO:0009409) and regulation of gene expression (GO:0010468) pathways were identified to be elevated in 8-oxoG depleted RNAs relative to background genes for both experimental conditions (Fig. 1I-J).

To provide a secondary validation of the enrichment status of the 8-oxoG-RIP-Seq identified RNAs, we performed an analogous enrichment technique under the same experimental stress conditions using the recently published ChLoRox-Seq method (26). This method relies on selective chemical labeling of 8-oxoG sites with a biotinylated linker molecule, subsequently enabling streptavidin mediated pulldown and next generation sequencing of 8-oxoG enriched RNAs. Encouragingly, we successfully validated several high confidence regions of elevated 8-oxoG accumulation (Supplemental Fig. S3A-B) (Table 1). The increased resolution of ChLoRox-Seq (≤ 500 nucleotide peak windows) enabled a more granular view of the location of 8-oxoG accumulation within these RNAs (Supplemental Fig. S4A-C). We did not observe any apparent locational bias of 8-oxoG enrichment within RNA transcripts (Supplemental Fig. S4D).

**Table 1.**
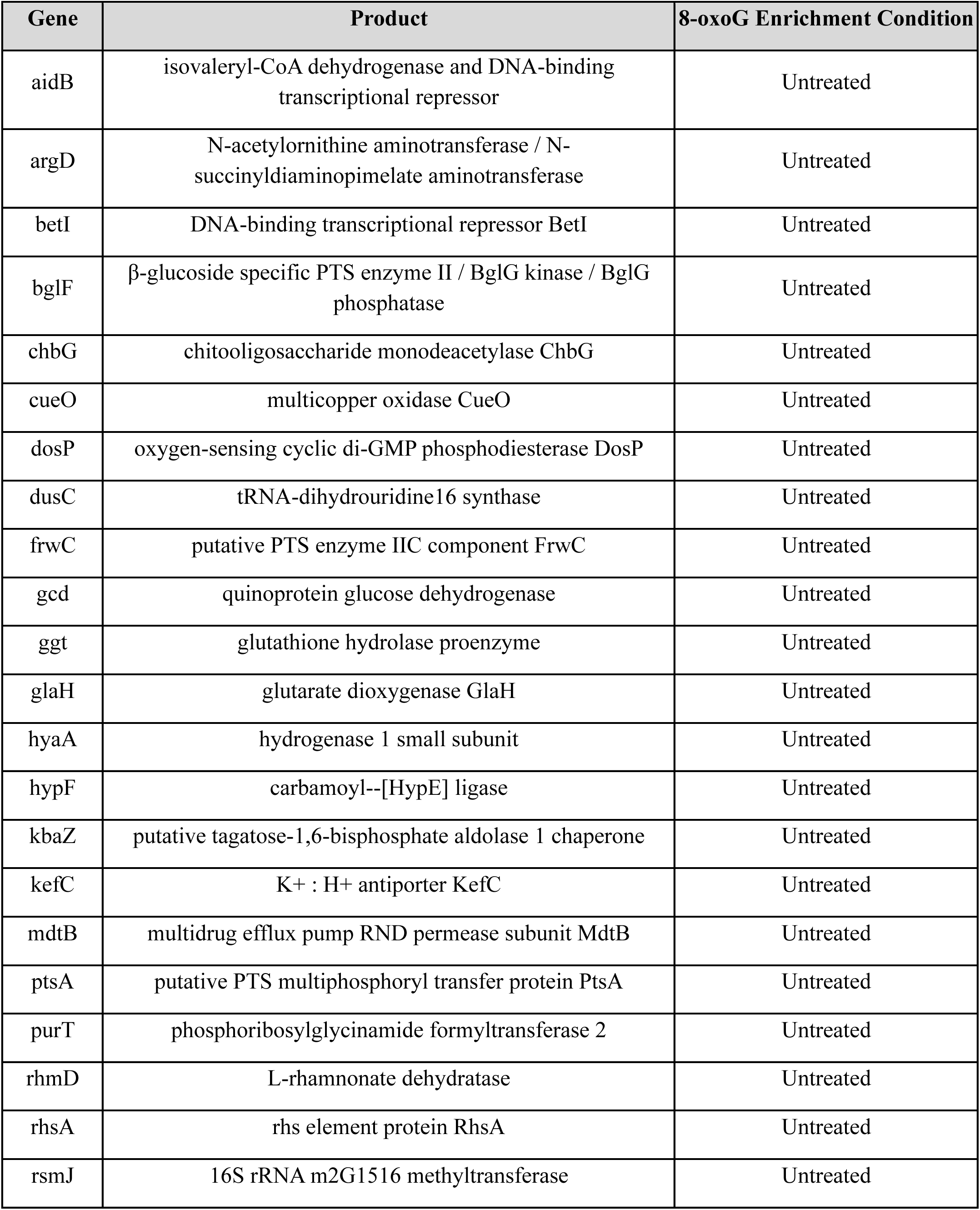
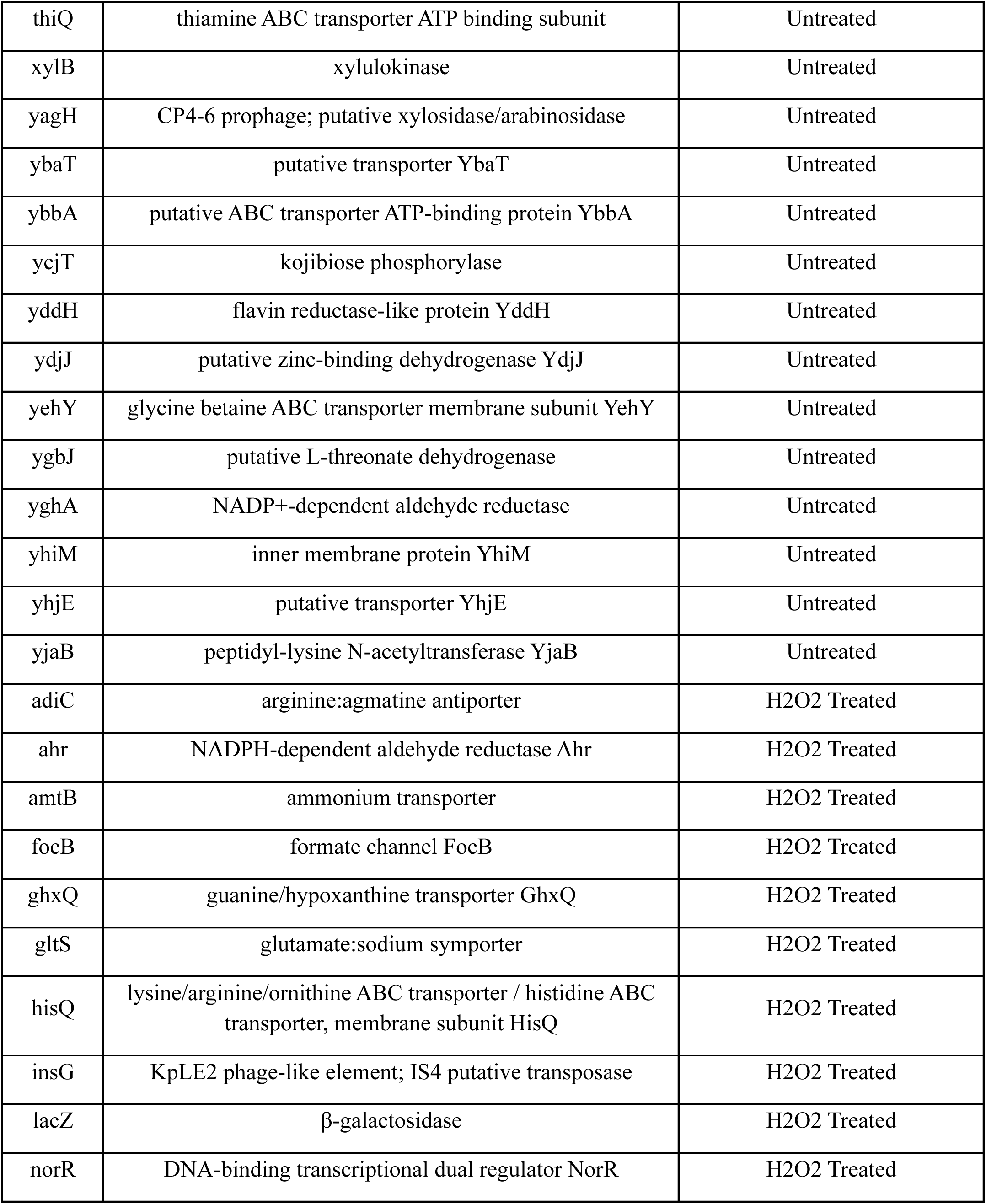
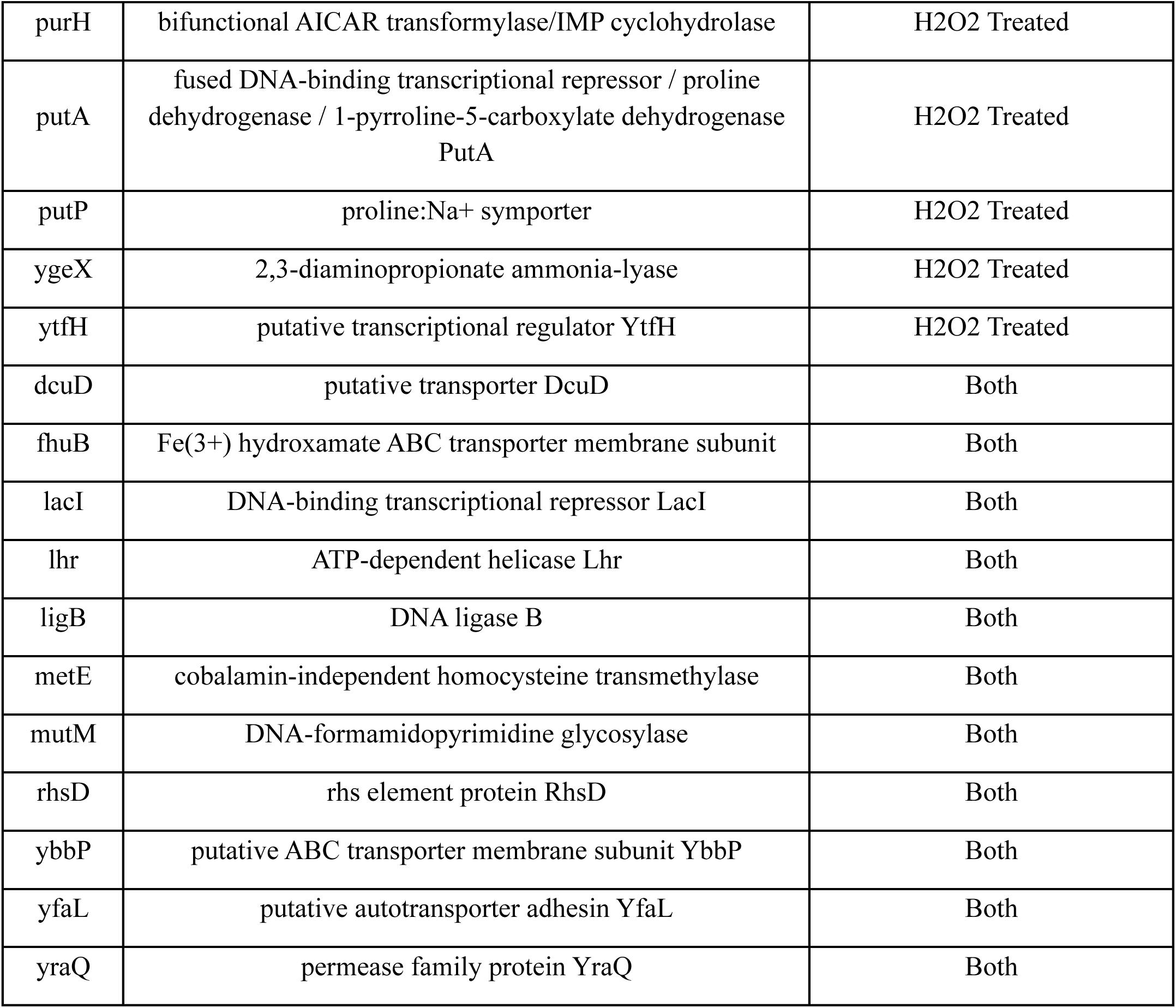
RNAs enriched in 8-oxoG modifications in *E. coli* identified by both 8-oxoG-RIP-Seq and ChLoRox-Seq detection methods across growth conditions.

| Gene | Product | 8-oxoG Enrichment Condition |
| --- | --- | --- |
| aidB | isovaleryl-CoA dehydrogenase and DNA-binding transcriptional repressor | Untreated |
| argD | N-acetylornithine aminotransferase / N-succinyldiaminopimelate aminotransferase | Untreated |
| betI | DNA-binding transcriptional repressor BetI | Untreated |
| bglF | β-glucoside specific PTS enzyme II / BglG kinase / BglG phosphatase | Untreated |
| chbG | chitooligosaccharide monodeacetylase ChbG | Untreated |
| cueO | multicopper oxidase CueO | Untreated |
| dosP | oxygen-sensing cyclic di-GMP phosphodiesterase DosP | Untreated |
| dusC | tRNA-dihydrouridine16 synthase | Untreated |
| frwC | putative PTS enzyme IIC component FrwC | Untreated |
| gcd | quinoprotein glucose dehydrogenase | Untreated |
| ggt | glutathione hydrolase proenzyme | Untreated |
| glaH | glutarate dioxygenase GlaH | Untreated |
| hyaA | hydrogenase 1 small subunit | Untreated |
| hypF | carbamoyl--[HypE] ligase | Untreated |
| kbaZ | putative tagatose-1,6-bisphosphate aldolase 1 chaperone | Untreated |
| kefC | K<sup>+</sup> : H<sup>+</sup> antiporter KefC | Untreated |
| mdtB | multidrug efflux pump RND permease subunit MdtB | Untreated |
| ptsA | putative PTS multiphosphoryl transfer protein PtsA | Untreated |
| purT | phosphoribosylglycinamide formyltransferase 2 | Untreated |
| rhmD | L-rhamnonate dehydratase | Untreated |
| rhsA | rhs element protein RhsA | Untreated |
| rsmJ | 16S rRNA m2G1516 methyltransferase | Untreated |
| thiQ | thiamine ABC transporter ATP binding subunit | Untreated |
| xylB | xylulokinase | Untreated |
| yagH | CP4-6 prophage; putative xylosidase/arabinosidase | Untreated |
| ybaT | putative transporter YbaT | Untreated |
| ybbA | putative ABC transporter ATP-binding protein YbbA | Untreated |
| ycjT | kojibiose phosphorylase | Untreated |
| yddH | flavin reductase-like protein YddH | Untreated |
| ydjJ | putative zinc-binding dehydrogenase YdjJ | Untreated |
| yehY | glycine betaine ABC transporter membrane subunit YehY | Untreated |
| ygbJ | putative L-threonate dehydrogenase | Untreated |
| yghA | NADP+-dependent aldehyde reductase | Untreated |
| yhiM | inner membrane protein YhiM | Untreated |
| yhjE | putative transporter YhjE | Untreated |
| yjaB | peptidyl-lysine N-acetyltransferase YjaB | Untreated |
| adiC | arginine:agmatine antiporter | H2O2 Treated |
| ahr | NADPH-dependent aldehyde reductase Ahr | H2O2 Treated |
| amtB | ammonium transporter | H2O2 Treated |
| focB | formate channel FocB | H2O2 Treated |
| ghxQ | guanine/hypoxanthine transporter GhxQ | H2O2 Treated |
| gltS | glutamate:sodium symporter | H2O2 Treated |
| hisQ | lysine/arginine/ornithine ABC transporter / histidine ABC transporter, membrane subunit HisQ | H2O2 Treated |
| insG | KpLE2 phage-like element; IS4 putative transposase | H2O2 Treated |
| lacZ | β-galactosidase | H2O2 Treated |
| norR | DNA-binding transcriptional dual regulator NorR | H2O2 Treated |
| purH | bifunctional AICAR transformylase/IMP cyclohydrolase | H2O2 Treated |
| putA | fused DNA-binding transcriptional repressor / proline dehydrogenase / 1-pyrroline-5-carboxylate dehydrogenase<br>PutA | H2O2 Treated |
| putP | proline:Na<sup>+</sup> symporter | H2O2 Treated |
| ygeX | 2,3-diaminopropionate ammonia-lyase | H2O2 Treated |
| ytfH | putative transcriptional regulator YtfH | H2O2 Treated |
| dcuD | putative transporter DcuD | Both |
| fhuB | Fe(3+) hydroxamate ABC transporter membrane subunit | Both |
| lacI | DNA-binding transcriptional repressor LacI | Both |
| lhr | ATP-dependent helicase Lhr | Both |
| ligB | DNA ligase B | Both |
| metE | cobalamin-independent homocysteine transmethylase | Both |
| mutM | DNA-formamidopyrimidine glycosylase | Both |
| rhsD | rhs element protein RhsD | Both |
| ybbP | putative ABC transporter membrane subunit YbbP | Both |
| yfaL | putative autotransporter adhesin YfaL | Both |
| yraQ | permease family protein YraQ | Both |

### Investigation of intrinsic RNA characteristics uncovers correlations with RNA 8-oxoG enrichment levels

As no enzymatic writer of 8-oxoG modifications is known, we hypothesized that differential deposition of this modification is probabilistically driven and likely biased based on intrinsic RNA characteristics. Previous work from our lab in human lung BEAS-2B cells demonstrated that RNA regions with elevated 8-oxoG modifications tend to occur within RNAs elevated in nucleotide length and G nucleotide composition (26). In the present work, we extended this analysis using our 8-oxoG-RIP-Seq dataset to evaluate several additional intrinsic RNA variables that may impact RNA 8-oxoG enrichment: G_1-5_ k-mer frequency (G:GGGGG), CDS length (Length), halflife (Halflife), relative cellular abundance as measured by TPM (Abundance), secondary structure as predicted by ViennaRNA (54) (basepair_prob), and G-quadruplex density as predicted by G4Boost (55) (per_nt_G4) (Fig. 2). We utilized 8-oxoG enrichment data from our Untreated experimental 8-oxoG-RIP-Seq condition in this analysis, as we were interested in understanding any potential correlations between these characteristics and the enrichment of RNA 8-oxoG marks at a basal level. To more clearly visualize apparent trends between 8-oxoG enrichment level and each intrinsic RNA property, we discretely subdivided the log2FoldChange distribution of 8-oxoG enrichment into 10 bins of roughly equal size (∼394 genes) (Fig. 2A). We subsequently evaluated the distribution of each intrinsic RNA variable within each discretized bin (Fig. 2B-K). Through this analysis, we observed that many of these intrinsic properties appeared to trend with RNA 8-oxoG enrichment. The observed trends for G frequency, Length, and Abundance variables were similarly found to trend with elevated RNA 8-oxoG accumulation through ChLoRox-Seq analysis (Supplemental Fig. S5A-C). Interestingly, G frequency within RNA 8-oxoG peak regions identified by ChLoRox-Seq did not significantly differ when compared to G frequency across the entire transcript CDS (Supplemental Fig. S5D).

**Figure 2.**
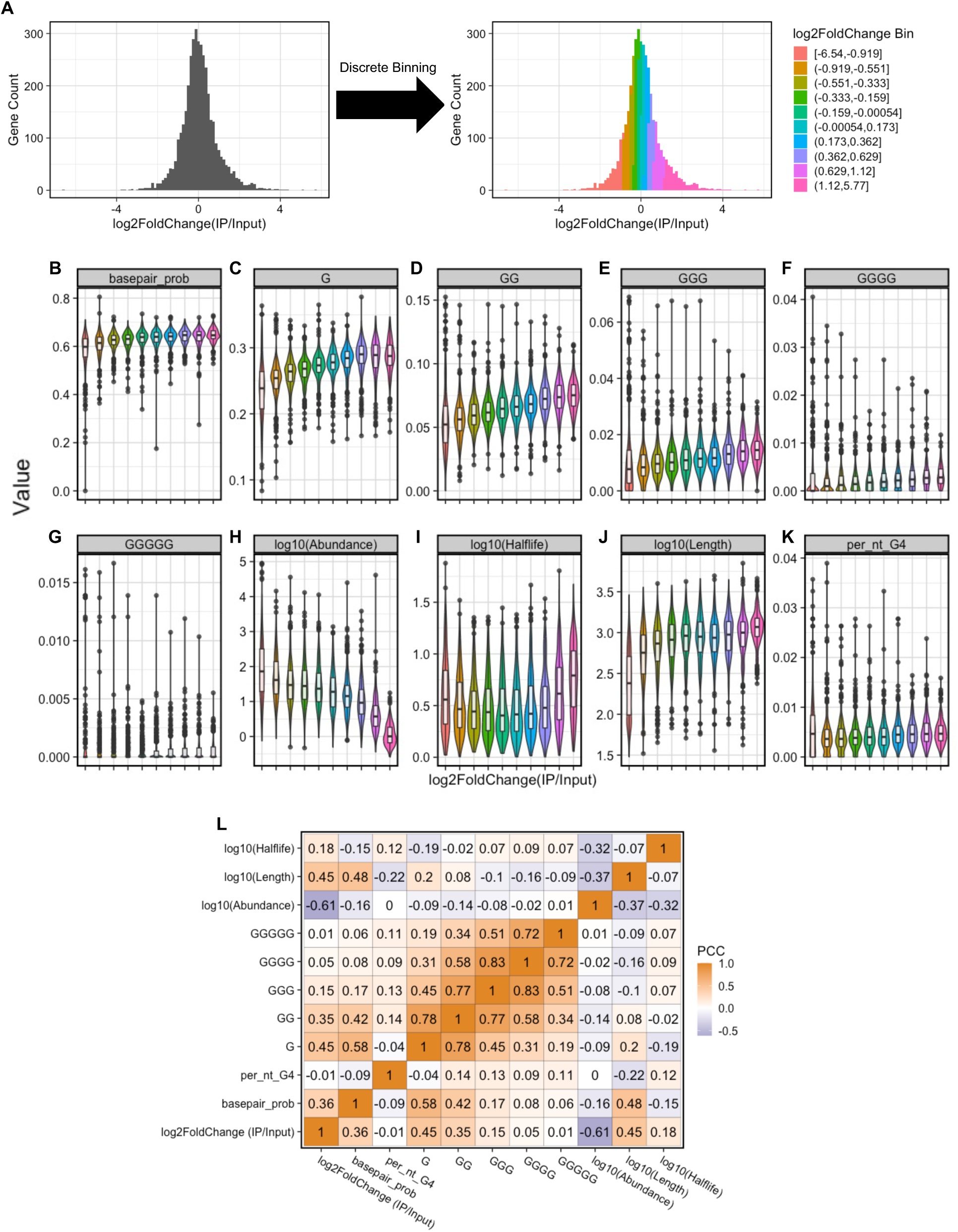
Intrinsic RNA properties correlate with levels of 8-oxoG enrichment. (A) Discrete binning approach used to evaluate trends between intrinsic variables and 8-oxoG-RIP-Seq-derived oxidation enrichment levels. (B-K) Boxplots overlaid onto violin plots for each discretized 8-oxoG enrichment bin measuring (B) secondary structure, (C) G frequency, (D) GG frequency, (E) GGG frequency, (F) GGGG frequency, (G) GGGGG frequency, (H) relative transcript abundance (TPM) (I) RNA halflife (min), (J) CDS length (nt), and (K) G-quadruplex density. (L) Correlation matrix of each intrinsic RNA property and oxidation enrichment levels. PCC: Pearson’s Correlation Coefficient.

To further evaluate trends amongst intrinsic RNA characteristics, we calculated the Pearson’s correlation coefficient (PCC) between each characteristic and 8-oxoG enrichment (Fig. 2L). This analysis further demonstrated the inherently correlated nature of these intrinsic RNA characteristics with 8-oxoG enrichment and identified several multi-colinear intrinsic variables (notably, the frequency of G_1-5_ k-mers). Our analysis revealed a modest positive correlation (*r* = 0.45) between RNA 8-oxoG enrichment and both G frequency and Length. Conversely, we observed a modest negative correlation (*r* = -0.61) between RNA 8-oxoG enrichment and Abundance. We found a slight positive correlation between RNA 8-oxoG enrichment and both basepair_prob and halflife (*r* = 0.36 and *r* = 0.18, respectively). Surprisingly, no apparent direct correlation between RNA 8-oxoG enrichment and per_nt_G4 was observed (*r* = 0.01). When considered in aggregate, these intrinsic characteristics were determined to likely influence the biased accumulation of 8-oxoG RNA modifications observed with the 8-oxoG-RIP-Seq assay.

### Multiple linear regression modeling predicts RNA 8-oxoG enrichment level given intrinsic characteristics

Based on our observation of the correlative nature of RNA 8-oxoG enrichment and intrinsic RNA characteristics, we sought to model these interactions (Fig. 3A). We chose a generalized multiple linear regression modeling approach to most clearly provide a mathematical link between these governing intrinsic RNA characteristics and 8-oxoG enrichment. We implemented 10-fold cross validation on a randomly selected 70% train set from our Untreated 8-oxoG-RIP-Seq dataset and simultaneously performed feature selection to only include parsimonious model variables. Based on our analysis, we determined that Abundance, Length, G mononucleotide frequency, GG dinucleotide frequency, basepair_prob, and per_nt_G4 were the most important variables for the model’s predictive performance (Fig. 3B). Our final model prediction achieved an adjusted coefficient of determination (*R^2^*) of 0.557 (Fig. 3C) on the 70% train set. We subsequently evaluated our model on the held out 30% test set, where it similarly achieved an adjusted *R^2^* of 0.571 (Fig. 3D), mean average error (MAE) of 0.425 (Fig. 3E), and mean ranked order distance of 164 (Fig. 3F). In general, our model underestimated the magnitude of enrichment for genes at the upper end of the 8-oxoG enrichment spectrum (Fig. 3C-D), ultimately yielding a more conservative estimate of 8-oxoG enrichment. Importantly, while our final trained model possessed a substantial MAE and modest adjusted *R^2^*, it captured predictive, statistically significant relationships between RNA 8-oxoG enrichment levels and intrinsic RNA characteristics (Table 2).

**Figure 3.**
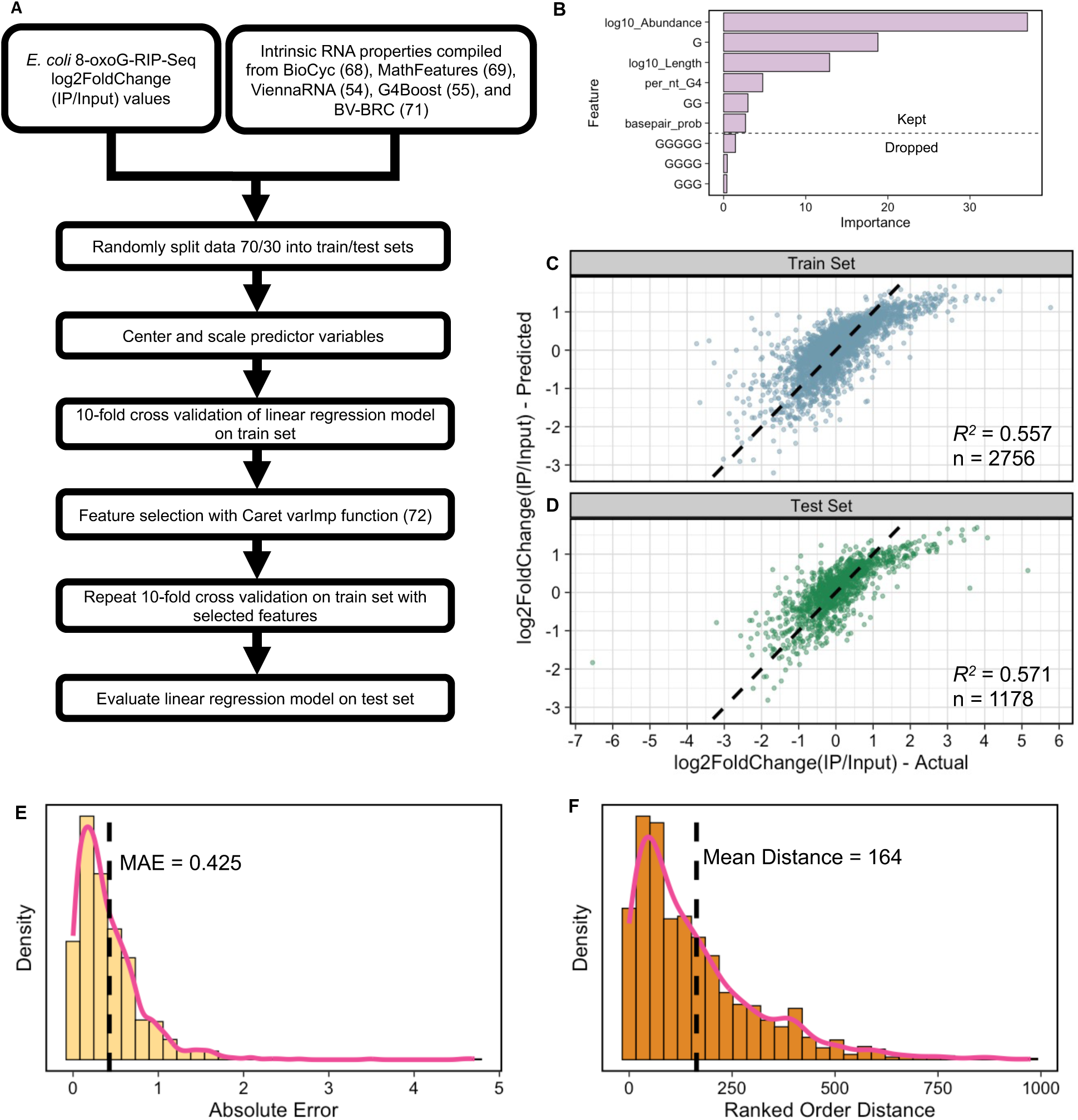
Multiple linear regression modeling predicts RNA 8-oxoG enrichment level in *E. coli* given intrinsic RNA properties. (A) Pipeline detailing approach used to build and evaluate multiple linear regression model. (B) Feature importance plot for each intrinsic variable used in building regression model. Variables above the dashed line were kept in the final model while variables below the dashed line were dropped from the final model. (C-D) Correlation plots showing regression model predicted versus actual 8-oxoG enrichment values for 70% train set (C) and 30% test set (D) Dashed lines represent y = x. (E) Histogram of absolute error for the regression model evaluated on the 30% test set. Vertical dashed line denotes the mean absolute error (MAE) Solid line represents a smoothed density fit to the histogram values. (F) Histogram of ranked order distance between regression model predicted 8-oxoG enrichment rank and actual oxidation enrichment rank. Vertical dashed line denotes the mean ranked order distance. Solid line represents a smoothed density fit to the histogram values.

**Table 2.** Final multiple linear regression model parameters for predicting 8-oxoG enrichment in *E. coli*.

| Variable | Coefficient | SE | t-value | p-value |
| --- | --- | --- | --- | --- |
| Intercept | 0.06082 | 0.01128 | 5.392 | 7.56e-08 |
| log10(Abundance) | -0.45654 | 0.01232 | -37.059 | < 2e-16 |
| G | 0.40730 | 0.02049 | 19.881 | < 2e-16 |
| log10(Length) | 0.18902 | 0.01423 | 13.285 | < 2e-16 |
| GG | -0.09361 | 0.01889 | -4.956 | 7.63e-07 |
| per_nt_G4 | 0.05763 | 0.01193 | 4.831 | 1.43e-06 |
| basepair_prob | -0.04188 | 0.01563 | -2.679 | 0.00742 |

### Multiple linear regression model predicts RNA 8-oxoG enrichment in diverse bacterial species, revealing a lack of conservation of 8-oxoG enrichment within major biological process pathways

Having developed a simplified, multiple linear regression model of 8-oxoG accumulation in *E. coli*, we were interested in further applying this model to a diverse repertoire of bacterial species. We hypothesized that the RNAs and, more generally, the biological process pathways predicted to be most enriched in RNA 8-oxoG modifications in *E. coli* might vary across bacteria. We selected *Deinococcus radiodurans* (*D. rad.*), *Mycolicibacterium smegmatis* (*M. smeg.*), *Candidatus Peligibacter ubique* (*Ca. P. ubique*), and *Shewanella oneidensis* (*S. one.*) for further application of our RNA 8-oxoG accumulation model based on their unique genome nucleotide compositions (high and low GC content) and their documented wide range of oxidative stress tolerances (12,13); furthermore, these species are of relevance to the broader scientific research community as model organisms. Upon applying our regression model to each of these species, we derived a predicted 8-oxoG enrichment level for each RNA. These predictions were converted to Z-scores within each species to generate a normal distribution of 8-oxoG enrichment. This post-processing step enabled a standardized evaluation of predicted RNAs with elevated propensity for 8-oxoG accumulation relative to the background gene population. Genes that were predicted to have an 8-oxoG enrichment level greater than one standard deviation (up or down) from the population mean were considered enriched or depleted in 8-oxoG modifications, respectively (Fig. 4A).

**Figure 4.**
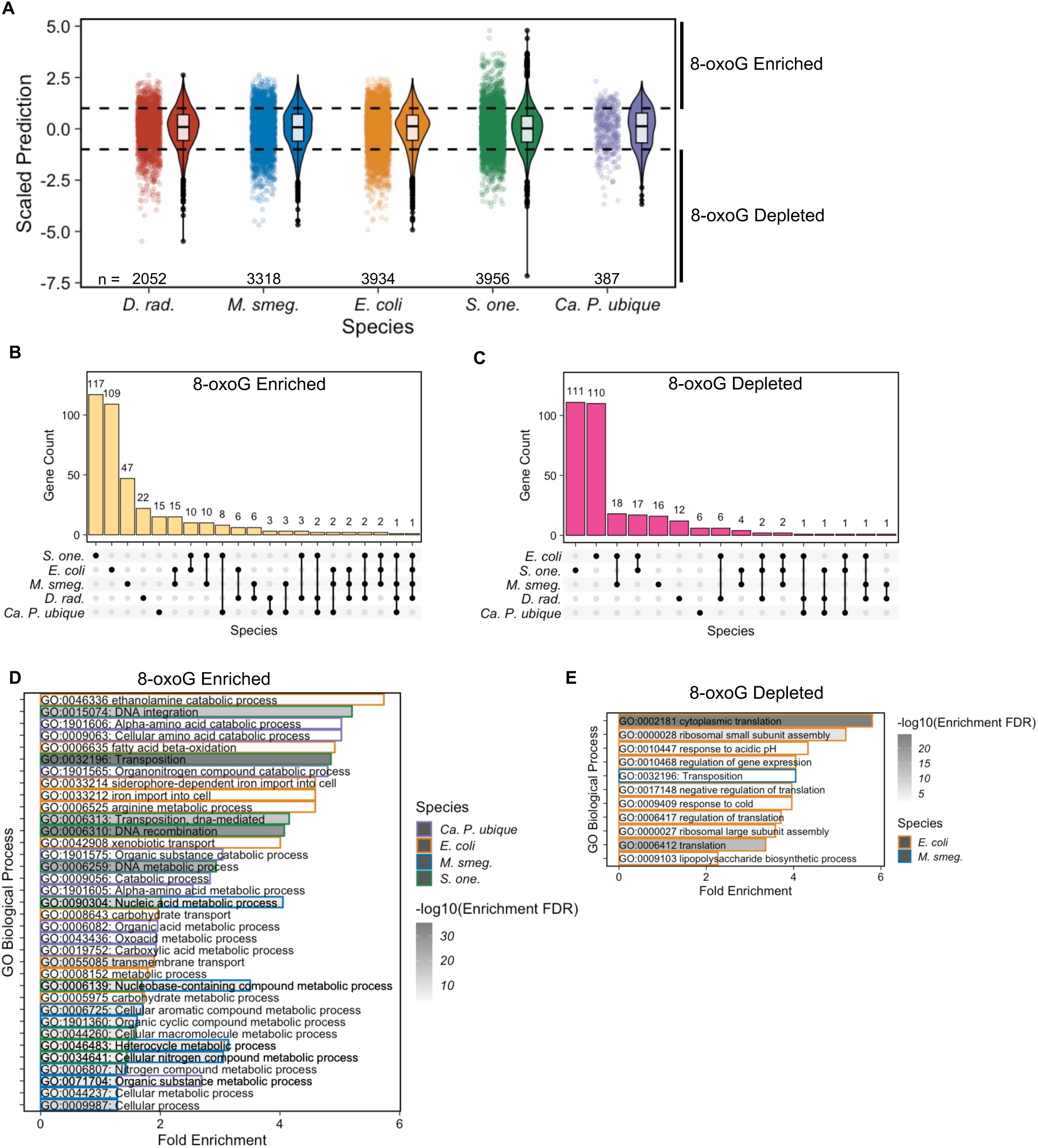
Application of linear regression model on diverse bacterial species predicts unique biological pathways with elevated RNA oxidation. (A) Boxplots overlaid onto violin plots beside jitter plots showing the distribution of scaled linear regression model predictions for each tested bacterial species. Horizontal dashed lines represent cutoffs for oxidation enriched (>1) and oxidation depleted (<-1) standard deviations away from the population mean. (B) Upset plot showing conservation of oxidation enriched genes across bacterial species. (C) Upset plot showing conservation of oxidation depleted genes across bacterial species. (D) Biological process gene ontology overrepresentation analysis for model predicted oxidation enriched RNAs across species. (E) Biological process gene ontology overrepresentation analysis for model predicted oxidation depleted RNAs across species.

We next investigated whether individual genes are predicted to be conserved in 8-oxoG enriched/depleted pools across bacterial species. To facilitate this analysis, we restricted the total gene pool to only include gene IDs that were shared in at least two of the modeled species. Our analysis revealed limited overlap between genes identified as naturally enriched or depleted in 8-oxoG across species (Fig. 4B-C). Of the 386 total genes identified as 8-oxoG enriched from all five modeled species, only 76 (∼20%) were shared between two or more organisms (Fig. 4B). Similarly, of the 309 total genes identified as 8-oxoG depleted, only 54 (∼17%) were shared between two or more species (Fig. 4C). While many of these predicted 8-oxoG enriched/depleted genes were unique to a single bacterial species, genes that are predicted across multiple species may point to conservation of 8-oxoG modification accumulation within targeted biological functions (Table 3).

**Table 3.**
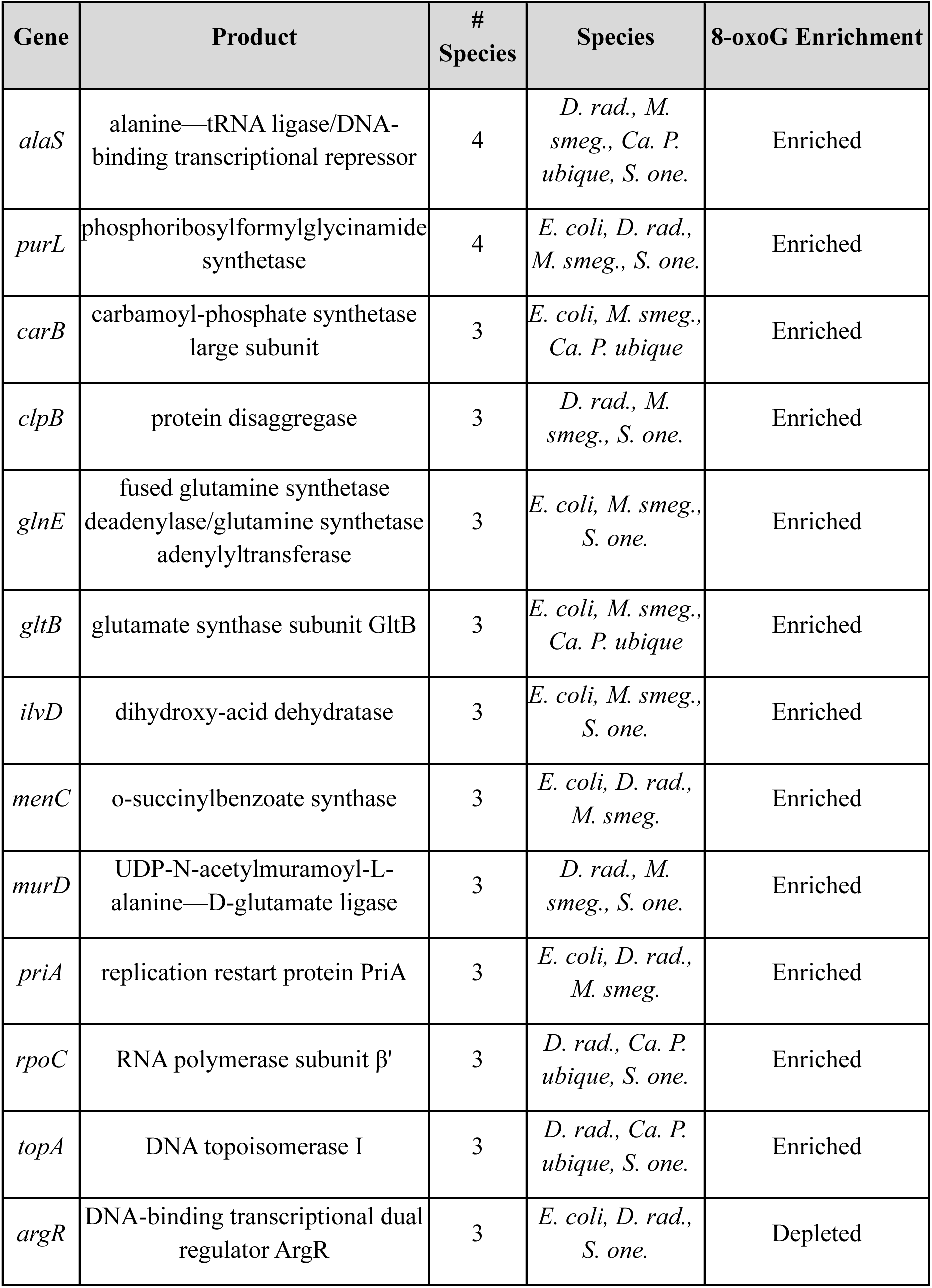
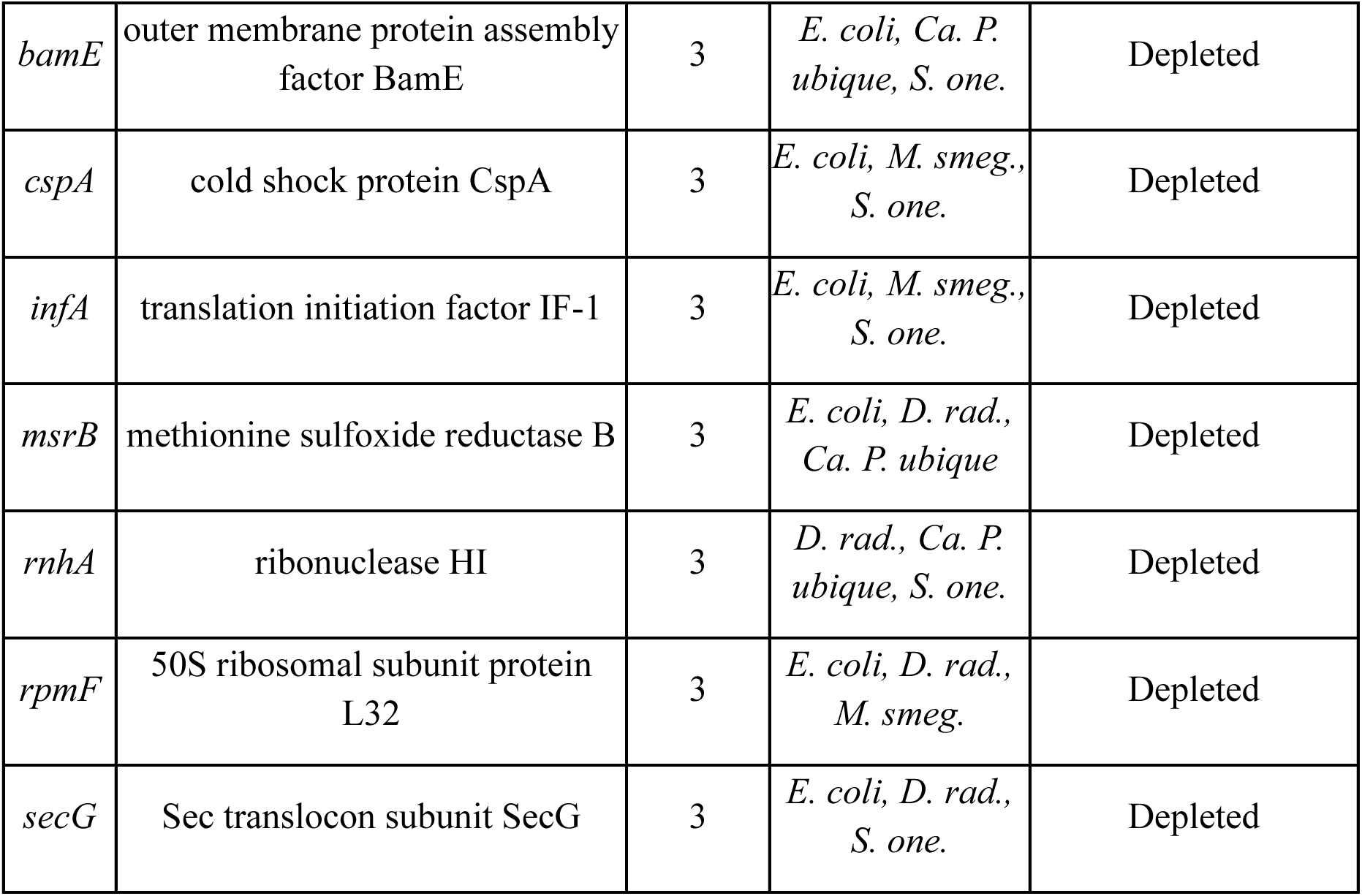
Genes predicted to be enriched/depleted in RNA 8-oxoG modifications across three or more of the modeled bacterial species.

We subsequently performed gene over representation analysis on 8-oxoG enriched RNA populations within each species to decipher whether certain biological processes were predicted to be conserved as elevated in 8-oxoG enriched or depleted RNAs across species (Fig. 4D-E). Encouragingly, gene over representation analysis of predicted 8-oxoG enriched and depleted RNAs in *E. coli* recapitulated many of the enriched GO pathways found from our 8-oxoG-RIP-Seq analysis (Fig. 4D-E, Fig. 1G-H). Most notably, we observed conservation of 8-oxoG enriched genes within pathways involved in transport processes (GO:0055085) and iron import (GO:0033212, GO:0033214) (Fig. 4D). Furthermore, evaluation of the predicted *E. coli* 8-oxoG depleted genes were similarly biased within pathways related to assorted translation processes (GO:0002181, GO:0006412) and response to cold (GO:0009409) (Fig. 4E). When we extended our gene over representation analysis to predicted 8-oxoG enriched/depleted genes in *D. rad*, we observed no significant enrichment (FDR < 0.05) within any biological process pathways. However, the nucleic acid metabolism pathway (GO:0090304) was found to be biased towards RNAs predicted to be enriched in 8-oxoG modifications in both *M. smeg.* and *S. one* (Fig. 4D). Interestingly, the transposition pathway (GO:0032196) was observed to be elevated in both predicted 8-oxoG *enriched* RNAs from *S. one.* and predicted 8-oxoG *depleted* RNAs from *M. smeg* (Fig. 4D-E). Overall, our modeling predictions indicated a low degree of overlap between GO biological process pathways elevated in 8-oxoG enriched/depleted RNAs across modeled species, concurrent with our prior finding regarding a lack of predicted conserved 8-oxoG enriched/depleted RNAs across species.

## Discussion

RNA oxidation remains understudied, particularly within the context of bacterial systems. In this work we performed, to our knowledge, the first transcriptome-wide analysis of the 8-oxoG modification enrichment landscape in *E. coli* (Fig. 1). From this analysis, we uncovered several unique intrinsic RNA characteristics that correlated with 8-oxoG modification enrichment (Fig. 2). Using these intrinsic characteristics, we trained and tested a multiple linear regression model to predict elevated accumulation of 8-oxoG marks within RNAs (Fig. 3). Although this model only showed modest predictive power, indicating its inability to completely replace the use of targeted sequencing experiments to truly understand these modification patterns, we captured significant variables that contribute to the biased accumulation of 8-oxoG marks (Table 2). We therefore applied this model to a small, diverse selection of four additional bacterial species to assess the degree of conservation of 8-oxoG accumulation landscapes within RNAs (Fig. 4). These chosen bacteria represented a wide range of oxidative stress tolerance, which we expected would display differences in their basal patterns of 8-oxoG accumulation. We found that the predicted patterns of 8-oxoG accumulation vary substantially across species, suggesting that molecular pathways potentially impacted by 8-oxoG accumulation are not widely conserved across bacteria.

### 8-oxoG modifications in E. coli unevenly accumulate within RNAs belonging to unique biological processes

Through leveraging an 8-oxoG-RIP-Seq approach, we first sought to identify RNAs that are both enriched and depleted in 8-oxoG modifications relative to the average of the entire population of RNAs across unstressed (Untreated) and oxidatively stressed (H_2_O_2_ Treated) conditions (Fig. 1). We were successful in capturing populations of 8-oxoG enriched (log_2_FoldChange(IP/Input) > 1, p_adj_ < 0.05) and depleted (log_2_FoldChange(IP/Input) < -1, p_adj_ < 0.05) RNAs under both experimental conditions (Fig. 1B-C). The disparate accumulation of 8-oxoG modifications across the *E. coli* transcriptome observed in this study suggests, as has been shown previously in eukaryotic systems (26–28,30), that certain RNAs within this bacterial cellular system are more prone to incurring oxidative lesions and, as a result, are subjected to altered RNA fates (e.g. degradation, translational errors, etc.) relative to RNAs with less 8-oxoG accumulation.

A key initial hypothesis underlying our work was that the landscape of 8oxoG-enriched RNAs would drastically shift under conditions of oxidative stress relative to basal conditions. However, our analysis revealed the opposite; a majority of the RNAs enriched in 8-oxoG under oxidative stress conditions were similarly enriched under unstressed conditions (283 of 383) (Fig 1D). Similarly, many of the RNAs depleted in 8-oxoG (149) were constant between oxidatively stressed and unstressed conditions (Fig 1E). Holistically, 8-oxoG enrichment levels were highly correlated between conditions (*r* = 0.90, Pearson’s correlation coefficient) (Fig 1F). Based on this analysis, we concluded that there is a significant level of basal 8-oxoG accumulation in *E. coli* RNAs and that the transcriptome-wide distribution of 8-oxoG marks is not as dependent on exogenous oxidative stress as we had initially postulated, at least under these defined experimental conditions. There remains a possibility that we failed to capture many oxidative stress induced regions of 8-oxoG accumulation due to the robustness of native ROS defense mechanisms, as we observed upregulation of several known ROS response elements under the tested 1 mM H_2_O_2_ stress condition (Supplemental Fig. S1A). Future 8-oxoG transcriptomic studies are necessary in *E. coli* which investigate additional oxidative stress sources, magnitudes, and exposure durations to enhance our understanding of how static the 8-oxoG modification landscape is under stress in this organism. Additionally, analogous future studies performed with genomic knockouts/knockdowns of key ROS response proteins may help uncover regions of 8-oxoG accumulation under stress that are normally critically protected.

We were next interested in investigating whether the populations of RNAs that were enriched/depleted in 8-oxoG modifications corresponded to specific functional pathways in the cell. Notably, through gene over representation analysis, we observed that the pool of 8-oxoG enriched RNAs contained a disproportionate number of transcripts encoding proteins involved in transport processes (Fig. 1 G-H). This observation might be partly due to intracellular RNA localization, as transporter proteins are frequently translated near the inner membrane through SRP translocation (56)—rendering their cognate RNAs less protected from membrane-permeable ROS diffusing into the cell from the outside environment. Consistent with this idea, previous work observed biased 8-oxoG accumulation within the FDFT1 transcript, which encodes a key protein for cholesterol biosynthesis, in human lung cells exposed to simulated air pollution mixtures (28). When we investigated biological pathways disproportionately represented within the 8-oxoG depleted RNA pool, we saw the emergence of many processes related to transcription, translation, and response to cold (Fig. 1 I-J). It is possible that these processes, being of key importance to cell survival, are inherently protected from oxidative modification. Overall, our findings from gene over representation analysis suggest that certain biological pathways and processes are disproportionately affected by the accumulation of 8-oxoG modifications within cognate RNAs in *E. coli*.

We next asked the question of whether specific RNA species engage in differential 8-oxoG accumulation under oxidative stress conditions, potentially connecting 8-oxoG modification enrichment to a larger cellular signaling cascade akin to the OxyR pathway at the protein level (8). We classified differentially modified RNAs as those possessing |log2FoldChange(IP/Input)_H2O2_ – log2FoldChange(IP/Input)_Untreated_| > 1.5. Our analysis revealed the emergence of several regulatory sRNAs within this differentially modified population (e.g. *istR*, *rttR*, *perR*, *fnrS*, *mgrR*, *sroA*, *dsrA*) (Supplemental Fig. S2). This result is intriguing, as the base-pairing promiscuity of 8-oxoG modifications to both A and C residues may significantly alter the targetome of these RNAs under conditions of oxidative stress and thereby contribute to cellular stress response through previously unforeseen mechanisms. Similar prior investigations into 8-oxoG in eukaryotic systems have uncovered differential modification of specific miRNA species within seed regions and subsequently connected modification status to an altered targetome (48–50). Additional follow-up studies are required to confirm and characterize the initial observations of differentially 8-oxoG modified sRNAs found in this work.

To validate initial findings from our 8-oxoG-RIP-Seq assay, we employed the use of an analogous pulldown and sequencing approach, ChLoRox-Seq (26), using the same experimental conditions. Although we did observe major differences in the populations of 8-oxoG enriched RNAs that were identified using this alternative method (Supplemental Fig. S3), perhaps due in part to the large variances observed across experimental replicates for each technique (Supplemental Fig. S6), we found several dozen RNAs that were consistently classified across methods (Table 1). Importantly, many of these shared RNAs encode for proteins involved in transport processes (e.g. *kefC*, *thiQ*, *ybaT*, *ybbA*, *yehY*, *yhjE*, *adiC*, *amtB*, *ghxQ*, *gltS*, *hisQ*, *putP*, *dcuD*, *fhuB*, *ybbP*, *yfaL*). It remains an open question whether the accumulation of 8-oxoG modifications within any of these specific RNAs plays a larger role in cellular response to oxidative stress by acting as sensors of ROS insult.

### Several intrinsic characteristics correlate with RNA 8-oxoG enrichment in E. coli

Based on findings from previous work in RNA and DNA which showed broad sequential and locational preferences for 8-oxoG accumulation sites (26,30,32,33), we hypothesized that 8-oxoG accumulation patterns in the *E. coli* transcriptome might be largely dictated by intrinsic RNA characteristics. We therefore investigated several RNA characteristics that we anticipated might be correlated with RNA 8-oxoG accumulation levels: G_1-5_ k-mer frequency, relative transcript abundance, CDS length, secondary structure, G quadruplex (G4) frequency, and halflife (Fig. 2). Our analysis of these properties revealed modest correlations between several intrinsic characteristics and 8-oxoG enrichment. Notably, relative abundance (Fig. 2H), CDS length (Fig. 2J), and G nucleotide frequency (Fig. 2C) appeared to most strongly bias the 8-oxoG enrichment level of RNAs. We validated the influence of these intrinsic RNA variables on RNA 8-oxoG enrichment by performing an analogous investigation on the ChLoRox-Seq dataset which confirmed similar trends (Supplemental Fig. S5A-C). Expectantly, we found a positive correlation between 8-oxoG enrichment and both G nucleotide frequency and CDS length, in line with previous findings (26). Interestingly, we found an inverse correlation between RNA 8-oxoG enrichment level and relative transcript abundance (Fig. 2H, L). This result was counterintuitive, as we anticipated higher relative intracellular abundance would probabilistically drive greater 8-oxoG enrichment levels. Instead, this result suggests that cells may partially combat oxidation of a given RNA by increasing expression levels. Importantly, these results supported our hypothesis that RNA 8-oxoG accumulation might largely be probabilistically driven, and that key intrinsic variables dictate which specific RNAs in the cell are more likely to be enriched or depleted in 8-oxoG.

### Accumulation of 8-oxoG in E. coli can be partially predicted through multiple linear regression modeling

Working off our findings that connected RNA 8-oxoG enrichment levels to intrinsic RNA characteristics, we subsequently exploited these relationships to construct a multiple linear regression model of RNA 8-oxoG enrichment (Fig. 3). This model demonstrated that intrinsic variables could be used to modestly predict relative RNA 8-oxoG enrichment, achieving an adjusted *R^2^* of 0.571 and MAE of 0.425 on the 30% held-out test set (Fig 3D-E). Importantly, despite high MAE, the model yielded a hierarchy of 8-oxoG enrichment that generally resembled the rank ordering of the 8-oxoG-RIP-Seq results (Fig. 3F). From these performance metrics, we grew confident towards using this regression model to make crude predictions of RNA 8-oxoG enrichment in additional, diverse bacterial species.

### Biological process pathways naturally enriched/depleted in 8-oxoG RNA modifications are not predicted to be widely conserved across diverse bacteria

Having constructed and validated our multiple linear regression model of RNA 8-oxoG enrichment in *E. coli*, we sought to apply this model to predict 8-oxoG enriched RNAs in four diverse bacterial species: *D. radiodurans, M. smegmatis, S. oneidensis,* and *Ca. P. ubique*. Our modeling approach identified populations of 8-oxoG-enriched and depleted RNAs within each species (Fig. 4A); although, the 8-oxoG enrichment status for these RNAs remains to be experimentally validated. Encouragingly, when we first applied our model to the *E. coli* transcriptome, we successfully predicted many of the same 8-oxoG-RIP-Seq identified, transport-related GO biological process pathways as being disproportionately elevated in 8-oxoG enriched RNAs (Fig. 4B, Fig. 1G). Similarly, in this organism, our model predicted pathways related to cold shock response and translation processes as being disproportionately elevated in RNAs naturally depleted in 8-oxoG modifications (Fig. 4C, Fig. 1I).

Assuming modest accuracy of our predictions, we next investigated whether the bias in functional pathways for proteins encoded by 8-oxoG-enriched/depleted RNAs that we observed in *E. coli* was consistent in other modeled bacteria. Our analysis into 8-oxoG enrichment predictions for the remaining modeled species revealed limited cross-species conservation of RNA 8-oxoG accumulation patterns (Fig. 4D-E). In *S. one.*, transposition-related pathways were predicted to be uniquely elevated in 8-oxoG enriched RNAs relative to the general RNA pool (Fig. 4D). This result suggests a possibility that this species relies on transposable elements to act as oxidation sinks, thereby protecting other cellular RNAs from incurring elevated oxidative damage. A similar phenomenon was recently suggested for the *ROSALIND* lncRNA as an ROS scavenger to protect mitochondrial translation machinery in human cells (30). Interestingly, these transposable elements are not predicted to be enriched in 8-oxoG for any of the other investigated species (Fig. 4D). In fact, the transposition pathway (GO:0032196) is predicted to be elevated in 8-oxoG *depleted* RNAs in *M. smeg.* (Fig. 4E). While the majority of GO biological process pathways elevated in 8-oxoG enriched/depleted RNAs were unique to a specific species, we observed that *S. one.* and *M. smeg.* shared a handful of 8-oxoG enriched pathways related to nucleic acid metabolism (GO:0090304, GO:0006139) and other metabolic pathways (GO:0046483, GO:0034641).

When we refined our focus of RNA 8-oxoG enrichment predictions to only include gene IDs that were detected in multiple (≥ 2) of the modeled species, we observed limited conservation of predicted 8-oxoG enriched and depleted RNAs (Fig. 4B-C) (Table 3). This low degree of gene overlap reinforces the idea that, as a non-enzymatically installed modification, 8-oxoG may be biased towards different transcripts (and thereby different biological pathways) across bacterial species. That is, it is possible that different bacterial species have evolved specific populations of RNAs to have intrinsic characteristics that are more or less susceptible to oxidations (such as those discussed above). However, it is worth noting that our simple regression model does not account for many species-specific biological factors (e.g. protein occupancy, nucleoid condensation, intracellular ion concentrations, etc.) that may have an outsized influence on the landscape of 8-oxoG modifications. Therefore, additional species-specific experimental investigations akin to the present study in *E. coli* are required to draw any definitive conclusions regarding the conservation of 8-oxoG landscapes across bacteria.

## Conclusion

Overall, our work into investigating transcriptome-wide 8-oxoG patterns in the model bacterium *E. coli* uncovered unique intrinsic RNA characteristics which appear to partially dictate RNA 8-oxoG accumulation. Using these characteristics, we successfully constructed a crude multiple linear regression model to predict RNA 8-oxoG enrichment in four other diverse bacterial species. The results from our model predictions suggest that RNA 8-oxoG modifications do not accumulate within conserved biological pathways across bacteria. Instead, we predict that the RNAs most enriched or depleted in 8-oxoG modifications can accumulate within specific biological pathways, but that these pathways are species dependent. Moving forward, the simple regression model created in this study can be used as a starting point for researchers interested in understanding the relative accumulation of RNA 8-oxoG marks in any bacterial species of interest.

### Limitations

The main limitation of our work on bacterial RNA 8-oxoG accumulation resides within a limited sample size and lack of targeted biochemical confirmation of true positive 8-oxoG sites. Our model reveals general trends in transcriptome-wide 8-oxoG accumulation; however, as evidenced by substantial MAE and modest *R^2^* (Fig. 3), we are unable to explain a considerable amount of the variance observed amongst RNAs possessing relative enriched/depleted 8-oxoG accumulation, possibly pointing to the influence of additional, uncharacterized biological variables (e.g. relative protein occupancy (57)). This result underscores the importance of continuing to perform transcriptome-wide investigations of RNA 8-oxoG enrichment using different next generation sequencing-based techniques (26–28). In this work, we have made initial predictions of the most 8-oxoG enriched and depleted RNAs in a small, diverse repertoire of bacterial species; however, these predictions remain to be biochemically confirmed through orthogonal techniques. As a result, the broad applicability of our model to study RNA 8-oxoG accumulation in other bacterial species remains unverified. Nevertheless, we have confidence that these trends should hold, as we have previously observed similar trends with respect to RNA length and G nucleotide frequency in human lung cells (26).

## Materials and Methods

### Cell culture

*Escherichia coli* str. K12 substr. MG1655 (Barrick Lab, UT Austin) was cultured aerobically in Luria-Bertani (LB) broth, shaking at 37°C and on LB-agar plates when required.

### Oxidative stress exposure treatment

*E. coli* cultures grown to mid-exponential phase (OD_600_ ∼0.5-0.7) were treated with either freshly diluted H_2_O_2_ solution to a final concentration of 1 mM (H_2_O_2_ stressed condition) or equal volume PBS (Unstressed condition). An incubation period of 20 minutes at 37°C was performed while shaking, followed by centrifugation to collect cell pellets. Pellets were immediately snap frozen in liquid nitrogen and held at -80°C until further use in RNA extractions.

### 8-oxoG RNA immunoprecipitation

RNA was extracted from cell pellets by the Trizol method and purified using the DirectZol RNA extraction kit following the manufacturer’s protocol. Elution was performed with N_2_-purged water to reduce artificial oxidation events. After extraction, samples were treated with DNase I (NEB) and repurified using the monarch RNA clean and concentrator kit. To remove ribosomal RNA, 9 μg of total RNA per sample were subjected to the Illumina rRNA depletion kit. Successful depletion and quality of samples was asserted through analysis by an Agilent 2100 Analyzer using the nano RNA assay. All intermediate buffers were freshly prepared and N_2_ purged the day of the experiment. A small portion of the depleted RNA was saved as the input fraction, and the rest was subjected to the RNA immunoprecipitation (RIP) protocol. The RIP was performed as previously described (27,28). Briefly, each RNA sample was incubated with 12.5 μg of 8-oxo-7,8-dihydroguanosine (8-oxoG) monoclonal antibody (clone 15A3, Bio-techne) for two hours at 4°C on IP buffer (10 mM Tris pH 7.4, 150 mM NaCl, 0.1% IGEPAL, and 200 units/ml SUPERaseIn RNA inhibitor). SureBeads Protein A magnetic beads (Biorad) were washed twice with 1X IP buffer and blocked by incubating for two hours at room temperature with BSA (2.5 mg/mL). After blocking, beads were washed twice with IP buffer, and a final resuspension was performed in IP buffer. Then, beads were incubated with the RNA-antibody samples rotating for two hours at 4°C. Following incubation, bead samples were washed three times (5 mins/wash) with 1X IP buffer. RNA was eluted by incubating samples with elution buffer (108 µg of 8-oxo-dG (Cayman Chemical) in IP buffer) for 1h at 4 °C. Elution samples were then cleaned using the Zymo RNA clean & concentrator kit. Input and IP samples were submitted to the UT Austin Genomic and Sequencing Analysis Facility (GSAF) for library preparation and RNA-seq. Libraries were prepared using the NEBNext® Ultra™ II Directional RNA Library Prep Kit for Illumina® and sequencing was performed on the NovaSeq 6000 instrument to obtain paired-end 150 length reads (10M reads per sample).

Raw sequencing .fastq files were processed as follows. Illumina sequencing adapters were trimmed using Cutadapt (version 5.0) (58) with the paired setting enabled and requiring a minimum quality score > 20. The trimmed files were checked for quality using fastqc (version 0.11.8) (59). The trimmed files were then aligned to the *E. coli* wild-type K-12 substr. MG1655 reference genome (GCF_000005845.2_ASM584v2_genomic.fna) using bwa mem (version 0.7.17) (60) with default parameters. Aligned SAM files were converted to BAM files, indexed, and sorted using samtools (version 1.20) (61). htseq-count (version 2.0.4) (62) was used to assign reads to gene features using the following parameters: -s reverse -m intersection-nonempty -t gene -i Name -a 30. Read count tables were filtered to only include genes with a mean read count >10 across all input fraction samples. Finally DE-Seq2 (63) was employed to evaluate regions of 8-oxoG enrichment (log2FoldChange(IP/Input)_Gene_ > 1, p_adj_ < 0.05) and depletion (log2FoldChange(IP/Input)_Gene_ < -1, p_adj_ < 0.05). All DE-Seq2 raw output tables are included in Supplemental Tables S1-S3.

### 8-oxoG RNA ELISA

RNA samples (2 μg) from H_2_O_2_ stressed or untreated conditions were subjected to Nuclease P1 (NEB, M0660S) digestion for 1 hr at 37°C. Samples were then dephosphorylated following the Quick-CIP protocol (NEB, M0525S). Bulk 8-oxoG levels from processed samples were directly measured using the DNA/RNA oxidative Damage (High Sensitivity) ELISA Kit (Cayman, 589320) following manufacturer instructions.

### ChLoRox-Seq

ChLoRox-Seq was performed in a similar fashion to previously published work (26) with the following slight alterations. RNA was extracted from snap-frozen cell pellets using a lysozyme assisted protocol. Briefly, pellets from were resuspended in freshly prepared lysozyme (Sigma) solution (2 mg/mL) in TE buffer and left incubating for 10 mins at RT. After incubation, DNA/RNA Shield Solution (Zymo Research, R1199-50) and DNA/RNA Lysis Buffer (Zymo Research, D7001-1-50) were added to the samples and mixed by pipetting. Proteinase K treatment was performed by incubating samples with the enzyme for 10 mins at RT. Then, samples were subjected to the Oligo Clean and Concentrator Kit (Zymo Research, D4061). RNA quality was assessed on an Agilent BioAnalyzer 2100; all samples generated a RIN score > 9.0. 10 µg of total RNA was depleted of contaminating rRNA species using the Illumina rRNA depletion kit (Illumina), following manufacturer protocol. The final elution step was performed using RNase-free, N_2_-purged H_2_O in lieu of the provided elution buffer. rRNA-depleted samples were fragmented using the NEB Magnesium RNA Fragmentation Module (NEB, E6150S) at 94°C for 4 minutes. Fragmented samples were re-purified using the Oligo Clean and Concentrator Kit (Zymo Research , D4061). Following purification, 8-oxoG modifications within RNA samples were labeled with biotin following the reaction conditions specified in (26) and again re-purified using the Oligo Clean and Concentrator Kit (Zymo Research, D4061). 1/10^th^ of each purified RNA sample was saved for use in creating the NGS input library. The remaining RNA was used in the Streptavidin Dynabead^TM^-mediated pulldown protocol performed in the identical fashion to (26). Pulldown RNA fractions were saved for use in creating the NGS pulldown libraries. Library preparation and sequencing was performed at the University of Texas at Austin Genomic Sequencing and Analysis Facility (GSAF). Libraries of all samples were prepared using the NEB Next Ultra II Direction RNA Library Prep Kit (NEB) following manufacturer protocol. Resulting libraries were tagged with unique dual indices and checked for quality and size using the Agilent High Sensitivity DNA Kit (Agilent). Library concentrations were measured using the KAPA SYBR Fast qPCR kit (Roche) and loaded for sequencing on a NovaSeq 6000 instrument (Illumina). Libraries were run in paired-end, 150 bp mode across two lanes to achieve a minimum sequencing depth of 20 million reads/sample.

Raw reads were processed as outlined above in *8-oxoG RNA Immunoprecipitation*. Aligned BAM files and the GCF_000005845.2_ASM584v2_genomic.gtf file were provided as input for the exomePeak2 analysis pipeline (64) to elucidate differential enrichment of reads around 8-oxoG modification sites. Regions of 8-oxoG enrichment were determined from the exomePeak2 program output by filtering for RPM_input_ > 0.5, log2FoldChange (Pulldown/Input) > 1, and FDR < 0.05. Raw outputs from the exomePeak2 pipeline are included in Supplemental Tables S8-S9.

### Statistics and data analysis

Data was visualized using the Tidyverse (65), ggsignif (66), and cowplot (67) packages in R. Unless otherwise specified, significance testing was performed using the Welch’s two-sided *t*-test.

### Curation of intrinsic RNA properties

#### RNA sequences

RNA CDS nucleotide sequences corresponding to the transcriptome of each bacterium were downloaded from the reference organism database on BioCyc (68). The MathFeature (69) kmer.py module was used to extract the k-mer frequencies from each RNA sequence.

#### Secondary structure

ViennaRNA (version 2.4.11) (54) was used for secondary structure prediction. The highest probability predicted structure (MFE) was selected after running the RNAfold algorithm on the full CDS of each gene. Base pairing probability was determined by taking the number of predicted base-paired nucleotides and dividing by gene nucleotide length.

#### G-quadruplex density

G-quadruplex (G4) predictions were made using the G4Boost algorithm (version 0.1.0) (55) with default parameters. Prediction probabilities of > 0.90 were classified as G4 hits. per_nt_G4 was calculated by normalizing the total number of predicted high probability G4 structures for a gene by its total length (nt).

#### Halflife

Halflife data for *E. coli* was curated from (70).

#### Relative Abundance

Raw gene read counts for non-*E. coli* bacteria were determined by importing RNA-sequencing data from relevant previously published work available on the sequence read archive (SRA) to the Bacterial and Viral Bioinformatics Resource Center (BV-BRC) RNA-Seq Analysis Tool (https://www.bv-brc.org/app/Rnaseq), utilizing the HT-Seq—DESeq2 analysis pipeline (71). A table with the specific reference genomes and SRA accession numbers used for each organism is provided in Supplemental Table S21. Raw read counts were filtered to exclude genes with a mean count >10 across 3 biological replicates. *E. coli* transcript abundances were taken from the count files derived from the bioinformatic processing steps outlined in *8-oxoG RNA Immunoprecipitation*. Relative transcript abundances (measured as transcripts per kilobase million (TPM)) for each gene were calculated using the following formula:

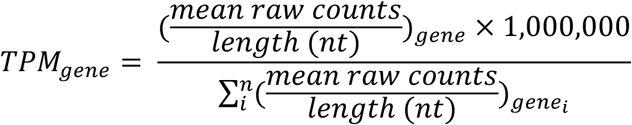

### Multiple Linear Regression Modeling

Our multiple linear regression modeling approach was carried out using the Caret package in R (72), following a conventional machine learning pipeline. In summary, the log2FoldChange 8-oxoG enrichment values determined for each gene in *E. coli* through 8-oxoG-RIP-Seq analysis (predicted variable) were randomly split 70/30 into train and test sets, yielding 2756 genes in the train set and 1178 genes in the test set. Intrinsic variable data for each gene were fed to the model during 10-fold cross-validation on the 70% train set. These intrinsic variables were centered and scaled to reduce model bias resulting from variable magnitude. Variable importance was evaluated using the Caret ‘varImp’ function to down-select variables for the most parsimonious model. 10-fold cross validation was again performed using the reduced variable set to create the final model. This model was then evaluated on the 30% held out test set. Final model parameters are included in Table 2.

Cross-species model predictions were made using the final version of the multiple linear regression model outlined above. Intrinsic properties for each bacterial species were fed into the model and output prediction values were converted to Z-scores to generate a normal distribution of 8-oxoG enrichment predictions. Predictions which scored greater than 1 standard deviation above the mean prediction value were classified as enriched in 8-oxoG. Predictions which scored greater than 1 standard deviation below the mean prediction value were classified as depleted in 8-oxoG. Complete species-specific intrinsic RNA metadata and model predictions are included in Supplemental Tables S10-S14.

### Gene ontology overrepresentation analysis

Genes passing specified significance/enrichment thresholding metrics were passed into the ShinyGO (version 0.82) webserver (https://bioinformatics.sdstate.edu/go/) with the appropriate reference organism selected (73). A background gene set was supplied which contained all genes detected in the respective analysis being performed. GO Biological Process enriched pathways were filtered to exclude pathways containing less than 9 total genes and an enrichment FDR < 0.05. Complete lists of significantly enriched biological process pathways are included in Supplemental Tables S4-S5, S15-S20.

## Data Availability Statement

Raw sequencing reads have been deposited as .fastq files in the NCBI Sequence Read Archive (SRA) database under BioProject ID PRJNA1345439. Any accessory data from this study will be shared by the corresponding author upon reasonable request.

## Author Contributions

This study was designed by M.R.B., B.Q.D., and L.M.C. Experiments and data analyses were performed by M.R.B and B.Q.D under the supervision of L.M.C. The manuscript was written by M.R.B. and edited by all authors. All authors have given approval towards the final version of the manuscript.

## Acknowledgements

The authors acknowledge the Texas Advanced Computing Center (TACC) at The University of Texas at Austin for providing high performance computing resources that have contributed to the research results reported within this paper. URL: http://www.tacc.utexas.edu. We also acknowledge the University of Texas Genomic Sequencing and Analysis Facility (GSAF) for library preparation and RNA sequencing. We thank members of the Contreras Lab for critical review of the manuscript. Lastly, the authors acknowledge the use of <u>Biorender.com</u> for graphical figure creation.

