## Supplemental Figures for "Profiling of RNA 8-oxoG marks in *Escherichia coli* identifies critical intrinsic characteristics that contribute to 8-oxoG accumulation in bacteria"

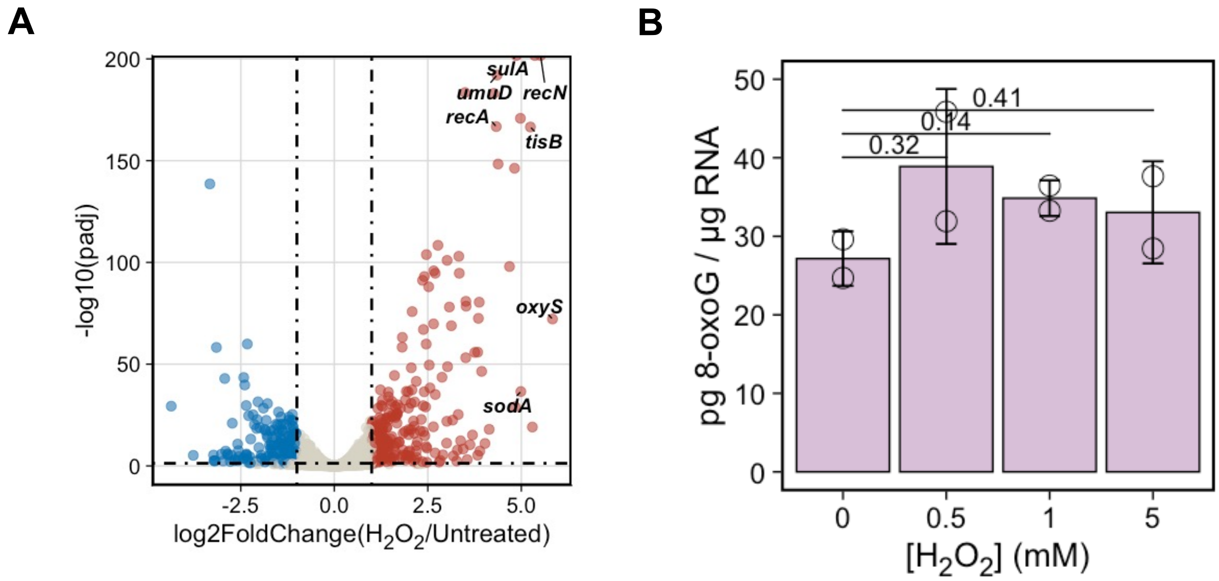


***Supplemental Figure S1.*** Oxidative stress conditions used in this study trigger a transcriptional response without increasing global 8-oxoG concentrations. (A) Volcano plot depicting differentially expressed genes between Untreated and $H_2O_2$ treated growth conditions. (B) Bar graph showing the amount of 8-oxoG (pg) per µg of total RNA input from RNA extracted from *E. coli* across a range of 20-minute $H_2O_2$ exposure concentrations, measured by ELISA. Circles represent individual biological replicates for each experimental condition ($n = 2$). No tested condition yielded statistically significant increases in free 8-oxoG concentration ($p > 0.05$, Welch's two-sided t-test).

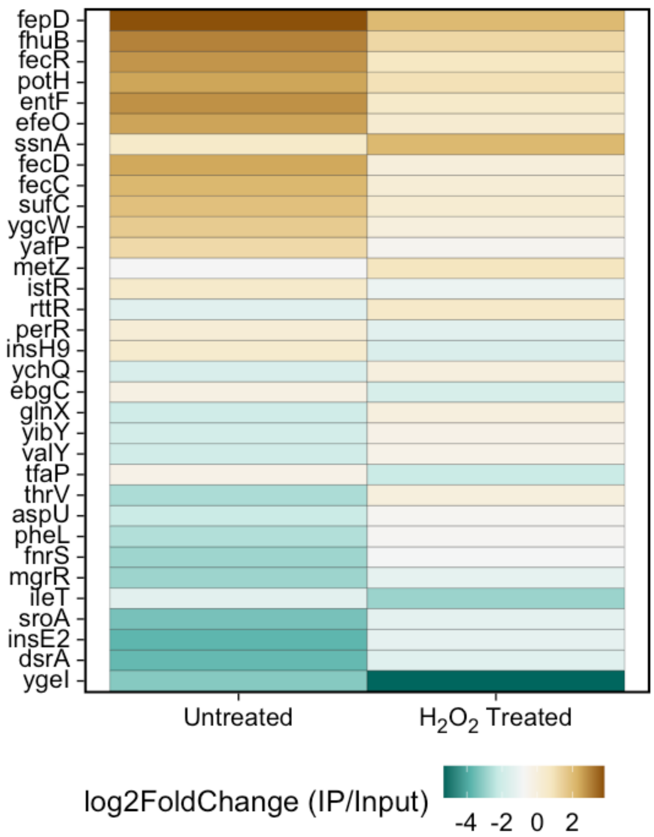


***Supplemental Figure S2.*** Differentially 8-oxoG modified RNAs ($|\text{log2FoldChange(IP/Input)}_{H2O2} - \text{log2FoldChange(IP/Input)}_{Untreated}| > 1.5$) between Untreated and $H_2O_2$ Treated conditions.

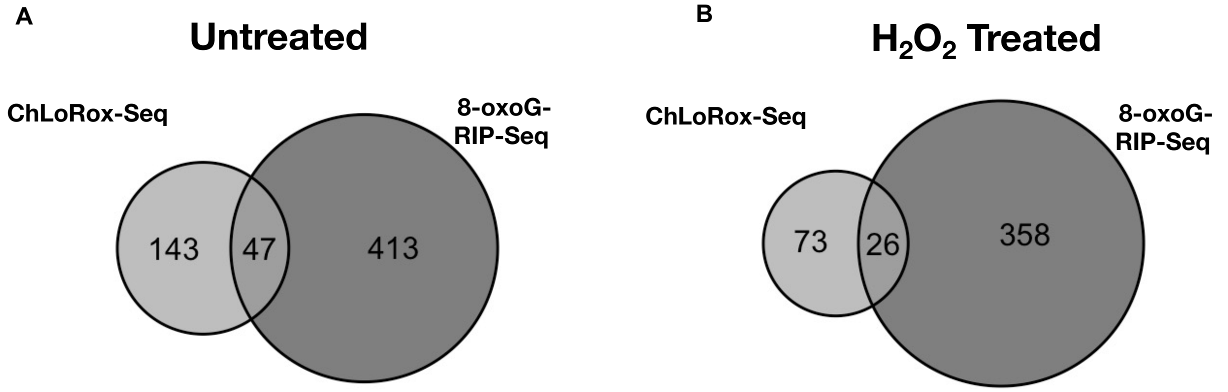


***Supplemental Figure S3.*** ChLoRox-Seq methodology recapitulates findings from 8-oxoG-RIP-Seq analysis. (A-B) Euler plots showing conservation of 8-oxoG enriched RNAs in Untreated (A) and $H_2O_2$ Treated (B) growth conditions across 8-oxoG-RIP-Seq and ChLoRox-Seq experiments.

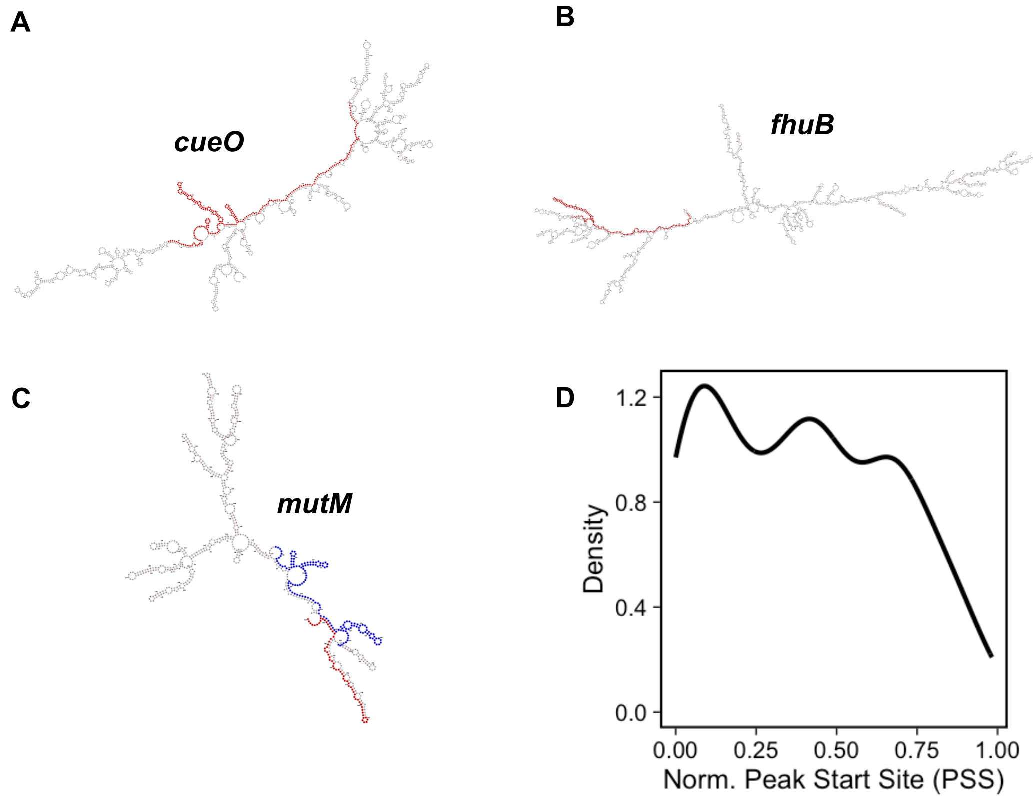


***Supplemental Figure S4.*** (A-C) ViennaRNA RNAfold predicted MFE secondary structure of representative conserved 8-oxoG enriched RNAs across both 8-oxoG-RIP-Seq and ChLoRox-Seq methodologies in the untreated condition ((A) *cuoE*, (B) *fhuB*, (C) *mutM*). Red/Blue coloring denotes specific region(s) of elevated 8-oxoG accumulation within RNAs identified by ChLoRox-Seq. (D) Density plot showing the distribution of 8-oxoG peak start sites (PSS) across transcripts normalized by transcript length.

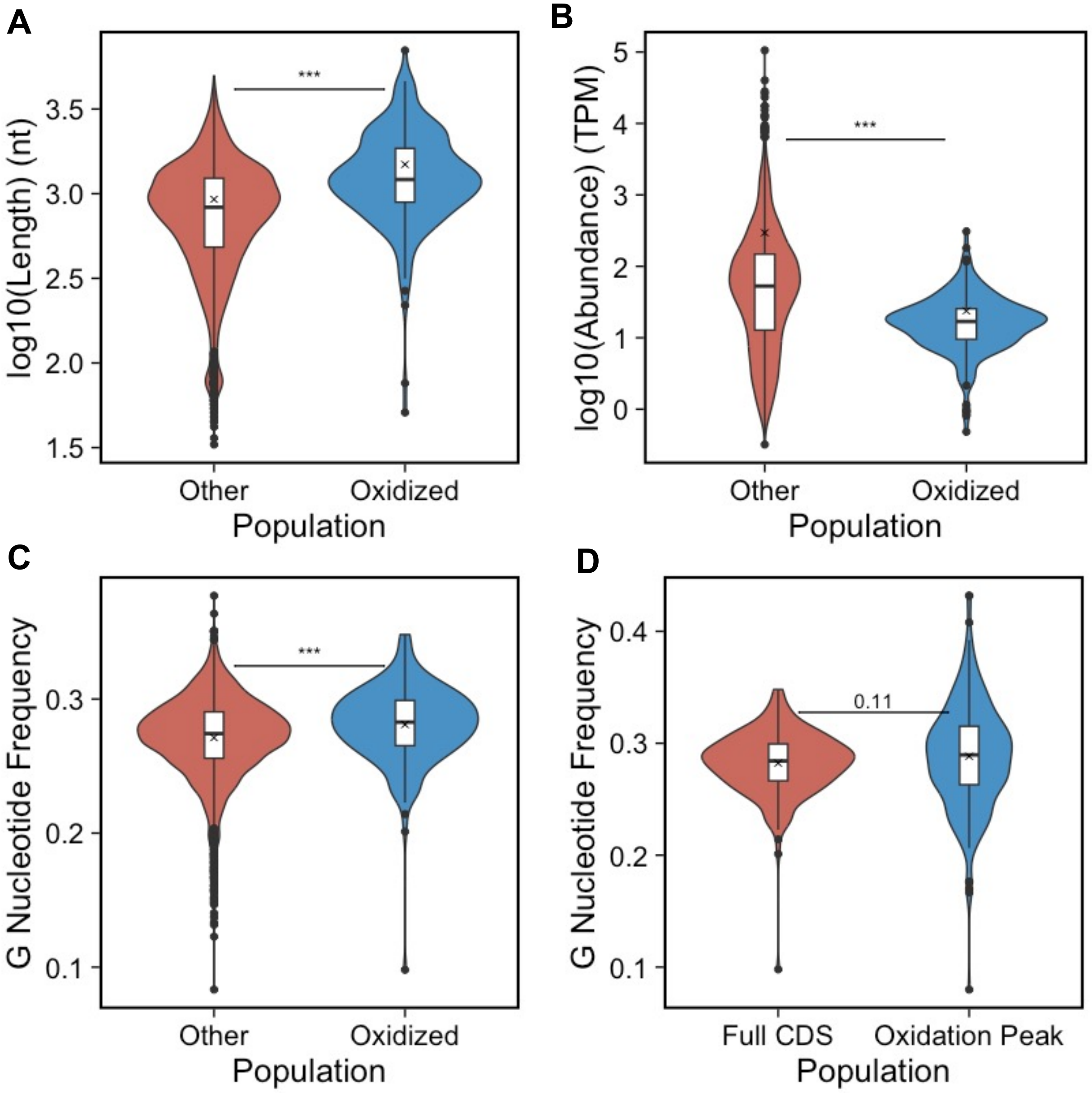


***Supplemental Figure S5.*** (A-C) Boxplots overlaid onto violin plots depicting the CDS length distributions (A), relative abundance distributions (B), and G nucleotide frequency distributions (C) of RNAs containing 8-oxoG peaks (Oxidized) versus all other RNAs detected in the Input fraction of ChLoRox-Seq experiment (Other). The "X" denotes population mean. (D) Boxplots overlaid onto violin plots depicting the G nucleotide frequency distributions of 8-oxoG peak regions (Oxidation Peak) versus the full CDS region (Full CDS) of RNAs harboring 8-oxoG peaks. The "X" denotes population mean. Statistical testing was performed using the unpaired Mann-Whitney U test. (*** $p<0.001$).

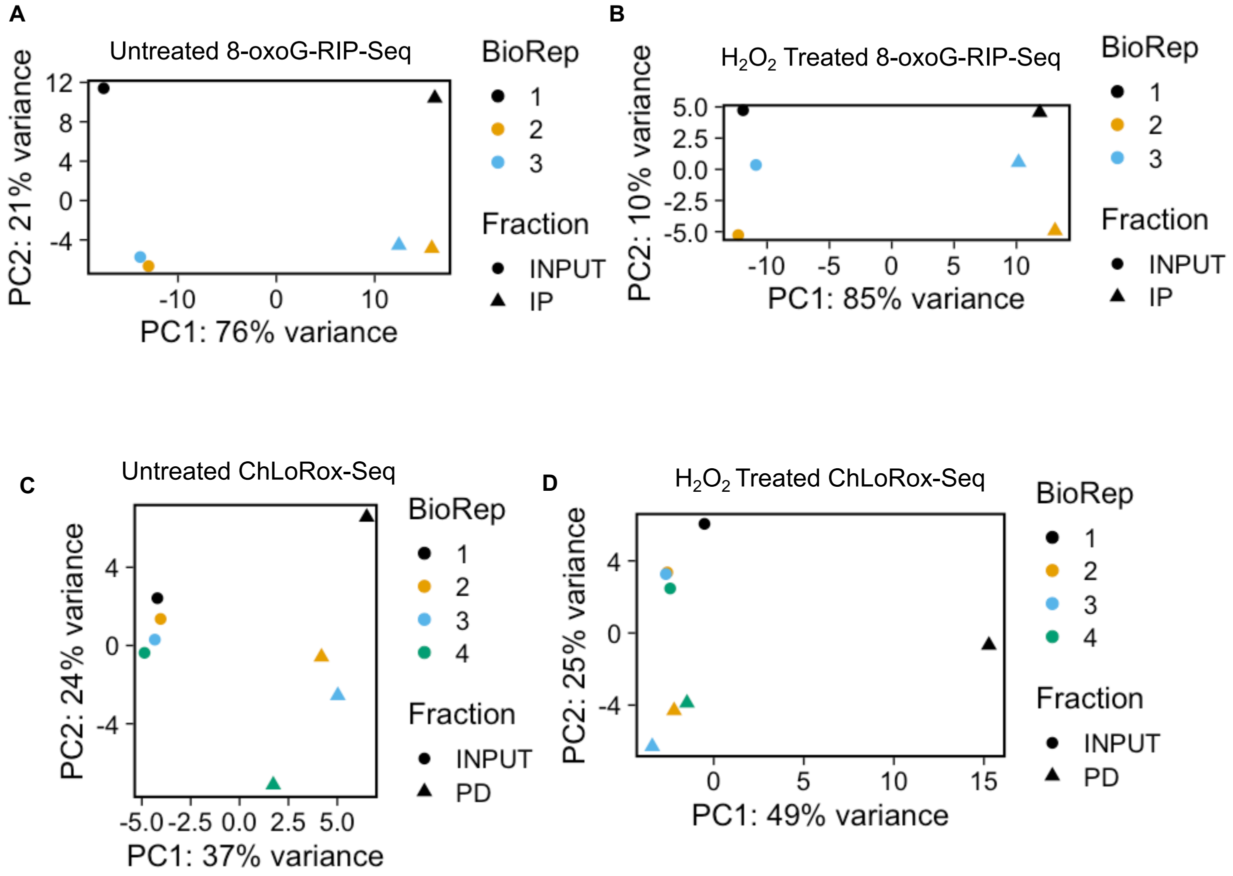


***Supplemental Figure S6.*** PCA plots of the top 500 gene feature counts for each biological replicate in pulldown experiments. (A-B) PCA plot of input (INPUT) and immunoprecipitated (IP) fractions of 8-oxoG-RIP-Seq experiment under Untreated (A) and H2O2 Treated (B) conditions. (C-D) PCA plot of input (INPUT) and pulldown (PD) fractions of ChLoRox-Seq experiment under Untreated (C) and H2O2 Treated (D) conditions.
